# PhysiBoSS 2.0: a sustainable integration of stochastic Boolean and agent-based modelling frameworks

**DOI:** 10.1101/2022.01.06.468363

**Authors:** Miguel Ponce-de-Leon, Arnau Montagud, Vincent Noël, Gerard Pradas, Annika Meert, Emmanuel Barillot, Laurence Calzone, Alfonso Valencia

## Abstract

Cancer progression is a complex phenomenon that spans multiple scales from molecular to cellular and intercellular. Simulations can be used to perturb the underlying mechanisms of those systems and to generate hypotheses on novel therapies. We present a new version of PhysiBoSS, a multiscale modelling framework designed to cover multiple temporal and spatial scales, that improves its integration with PhysiCell, decoupling the cell agent simulations with the internal Boolean model in an easy-to-maintain computational framework. PhysiBoSS 2.0 is a redesign and reimplementation of PhysiBoSS, conceived as an add-on that expands the PhysiCell agent-based functionalities with intracellular cell signalling using MaBoSS having a decoupled, maintainable and model-agnostic design. PhysiBoSS 2.0 successfully reproduces simulations reported in the former version and expands its functionalities such as using user-defined models and cells’ specifications, having mechanistic submodels of substrate internalisation with ODEs and enabling the study of drug synergies. PhysiBoSS 2.0 is open-source and publicly available on GitHub (https://github.com/PhysiBoSS/PhysiBoSS) under the BSD 3-clause license with several repositories of accompanying interoperable tools. Additionally, a nanoHUB tool has been set up to ease the use of PhysiBoSS 2.0 (https://nanohub.org/tools/pba4tnf/).

## Introduction

Cancer is a highly complex and heterogeneous disease caused by alterations in signalling pathways that control key cellular mechanisms as well as population dynamics [1–3]. Understanding cancer biology, drug treatments and relapse can be greatly improved by the use of mathematical models [4–6] that identify points of intervention. These can then be targeted using drugs with high efficacy and potency that increase the therapeutic effect of the cancer treatment [7, 8]. In fact, mathematical models can be used to identify combinatorial therapies that reduce drug resistances caused by the activation of compensatory cancer pathways in response to targeted therapies that only inhibit a single target [7, 9–11]. These combinatorial therapies can make use of synergies among drugs enabling lower drug dosages and, thus, reducing unwanted side effects while retaining the desired drug effects [12, 13].

Examples of such mathematical frameworks include Boolean modelling which can uncover clues in a cell’s intracellular signalling that lead to disease [11, 14] and agent-based modelling that can help understand the role of the interactions between cells and their environment [15–17]. Furthermore, these two frameworks can be integrated into multiscale simulations which can be used to explore the combined effect of genetic and environmental perturbations, as in the case of PhysiBoSS. PhysiBoSS [18] is a standalone multiscale simulation software that was implemented by coupling an early version of PhysiCell [15], a multiscale agent-based modelling framework that allows the simulation of populations of cells in a defined microenvironment, together with MaBoSS [19, 20], a continuous-time Markovian simulator for Boolean models that describe the cell’s intracellular signalling and regulatory networks [14]. The introduction of this hybrid simulation framework was an important step toward the mechanistic multi-scale description of complex biological systems such as healthy tissues and tumours [16]. Nevertheless, in its original version, PhysiBoSS presented some design problems. Specifically, PhysiBoSS 1.0 lacks a clear interface between these two tools which resulted in a software package that is hard to modify and extend without changing core functionalities, hindering its long-term maintenance and usability. Furthermore, it was designed and tested using a specific signalling model with a set of predefined phenotypes making it hard to adapt or extend to use different models and use cases.

Herein, we present PhysiBoSS 2.0, a new design and reimplementation of PhysiBoSS which solves the problems of its early version, extends its functionalities and enables a more flexible definition of models. PhysiBoSS 2.0 was implemented as an add-on component of PhysiCell that provides access to the MaBoSS simulator in a clear and transparent way. In this new design, both PhysiCell and MaBoSS are decoupled and therefore can be upgraded independently (Figure 1A). Furthermore, the code was designed and implemented following the best practices guidelines for bioinformatics software development, ensuring its long-term maintenance and prolonging its lifespan [21].

**Fig. 1.**
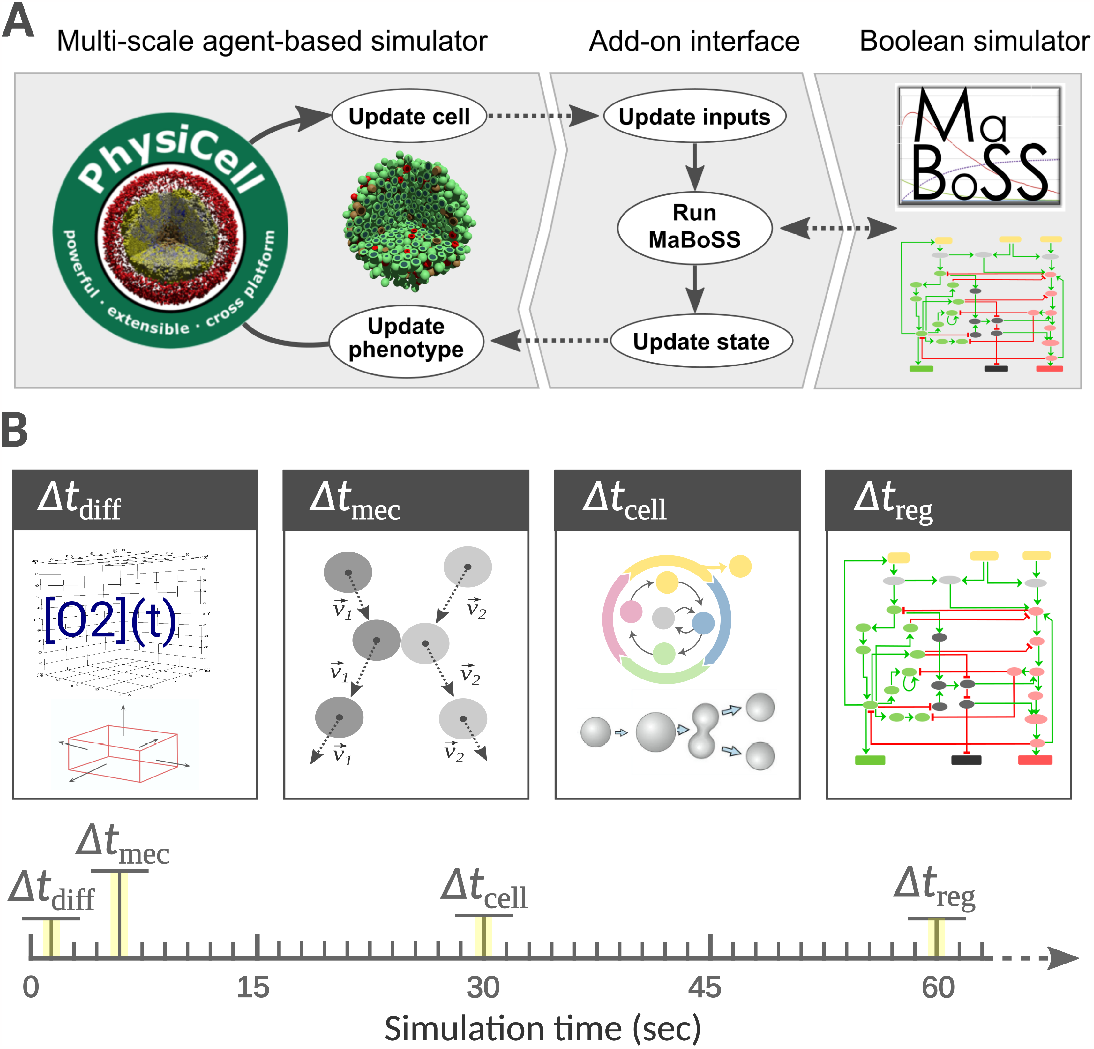
PhysiBoSS 2.0 add-on-based design. Panel A shows a diagram of the add-on-based design of PhysiBoSS 2.0 that decouples PhysiCell and MaBoSS providing Boolean simulation functionality to individual cell agents in a maintainable manner. Panel B depicts the different time scales used in PhysiBoSS simulations which include: the Δ*t*_*diff*_ time scale where diffusion, uptake and secretion processes are updated; Δ*t*_*mec*_ where the mechanics (movement and physical interactions) are updated; Δ*t*_*cell*_ in which cell processes such as volume, cell cycle and death models are updated; and Δ*t*_*reg*_ the regulatory time scale in which Boolean models are updated by running MaBoSS simulator.

PhysiBoSS 2.0 includes new functionalities like allowing the use of highly customisable settings such as user-defined Boolean models and cell types defined in the configuration XML, by easing the building of user-defined examples with different Boolean models in different cell types. In this work, we show different approaches for integrating Boolean models into multiscale simulations using the examples presented in the original publication. Moreover, we have used PhysiBoSS 2.0 new features to study the drug effects and synergies in multiscale simulations [22, 23] of a prostate cancer cell line, LNCaP, with six available drugs [24]. We have validated the single-drug treatments using experimental data and inspected closely the effect of drug dosages in the generation of heterogeneities in the cell population. Altogether, we show that PhysiBoSS 2.0 is a step towards the development of a PhysiCell add-on ecosystem of different types of models [25] that allows for the scaling-up of simulations in exascale high-performance computing clusters [22, 26, 27].

## Methods

### Boolean models

In this subsection, we describe the different Boolean models used in this work. The first model is focused on the effect of TNF presence on cell fate decisions, was used previously in PhysiBoSS 1.0 and is an extension of a published Boolean model of cellular fates [14]. It has 31 total nodes, 25 internal nodes, 3 input nodes (TNF, FADD, FASLG) and 3 output nodes (*Survival, Non-Apoptotic Cell Death* (NonACD), *Apoptosis*). The second model is a general model of prostate cancer that was used to identify potential drug targets in prostate cancer that were personalised to patients and cell lines using different *omic* datasets [24]. The model includes a total of 133 nodes of which 9 correspond to input nodes and 6 to outputs nodes. The personalized version of the model includes six prostate cancer cell lines (LNCaP, 22Rv1, BPH1, DU145, PC3 and VCaP), nevertheless, the results here presented were obtained using LNCaP. All the models are available in the dedicated repository: https://github.com/PhysiBoSS/Boolean-models.

### Personalisation of Boolean models

Boolean models can be used to simulate the effect of therapeutic interventions and predict the expected efficacy of candidate drugs on different genetic and environmental backgrounds by using our PROFILE v2 methodology [24]. Herein, the prostate Boolean model was tailored to different data sets using PROFILE v2 methodology to obtain personalised models that capture the particularities of a set of patients [28] and cell lines [29]. Proteomics, transcriptomics, mutations and copy number alteration (CNA) data can be used to modify different variables of the MaBoSS framework, such as node activity status, transition rates and initial conditions. The resulting ensemble of models is a set of personalised variants of the original model that can show great phenotypic differences. Different strategies (use of a given data type to modify a given MaBoSS variable) can be tested to find the combination that better correlates to a given clinical or otherwise descriptive data. In the present case, prostate-cell-line-specific models were built using mutations, CNA and RNA expression data. More details on PROFILE methodology can be found in its own work [28] and at its dedicated GitHub repository: https://github.com/PhysiBoSS/PROFILE_v2. Note that apart from using omics data to personalise the Boolean model, PhysiBoSS 2.0 can also include cell-line-specific phenotypic data, for instance doubling times, to tailor the simulations to a desired behaviour (see Section *Personalisation cell lines in PhysiBoSS 2*.*0* in Supplementary Text).

### Simulating drug inhibition synergies in the prostate model

In *PhysiBoSS 2*.*0*, diffusing drugs in the microenvironment target and inhibit specific nodes in the Boolean model within the individual cell agents. As modellers, the goal is to identify drugs that target candidate nodes and specific treatment strategies that together might be relevant in producing desired therapeutic outcomes such as increased apoptosis or reduced proliferation. To illustrate how *PhysiBoSS 2*.*0* can be used to conduct *in-silico* drug screening experiments, we selected a list of target nodes from [24] that have yielded interesting results using prostate Boolean models. Suitable drugs were then identified using the DrugBank database [30] and matched with their availability on the Genomics of Drug Sensitivity in Cancer (GDSC) database [31]. The drug-target pairs of interest for LNCaP can be found in Table 1.

**Table 1.**
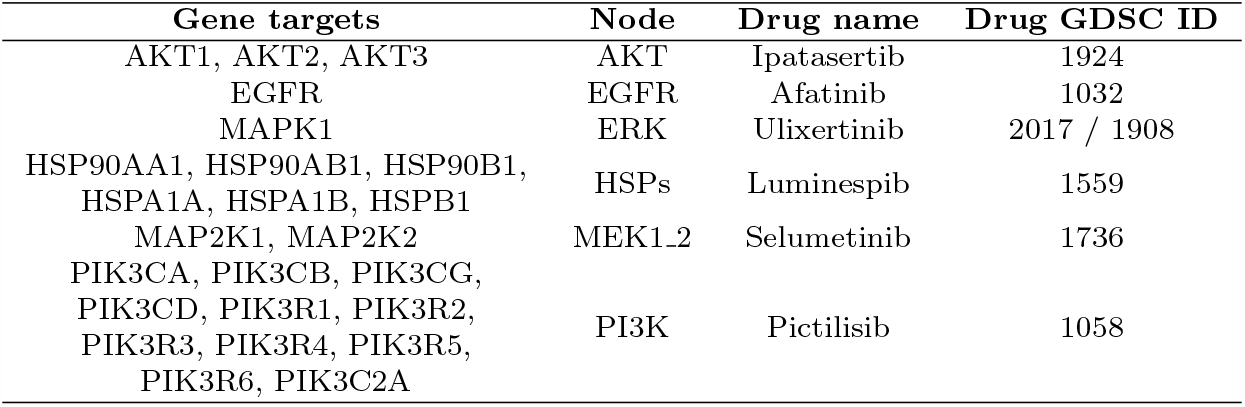
Drug-target pairs used to perform the drug simulations on the LNCaP-specific Boolean model.

We then integrate the estimated dose-response curves (see section *Estimation of drug-response for single and double drug studies*) into *PhysiBoSS 2*.*0* to model the effect that the local concentration of a drug in the nearby surroundings of a cell has on the intracellular signalling model. Specifically, a cell agent is informed of the drug concentration for each simulated substrate at its nearest voxel. Then, the corresponding target node is inhibited with a probability calculated based on the dose-response curve for the selected drugs (Table 1). The probability of inhibiting a node in a given cell is equal to 1 − *f* ([*X*]_*i*_) where [*X*]_*i*_ is the concentration of the drug in at voxel *i* and *f* is the normalised dose-response fitted function.

We simulated the inhibition of six nodes of interest on the LNCaP model by using *PhysiBoSS. MaBoSS* simulation engine, integrated into *PhysiBoSS*, can perform simulations changing the proportion of activated and inhibited status of a given node. For instance, out of 5000 trajectories of the Gillespie algorithm, *MaBoSS* can simulate 70% of them with an activated AKT and 30% with an inhibited AKT node. Then, the phenotypes’ probabilities for the 5000 trajectories are averaged, and these are considered to be representative of a model with a drug that reduces the activity of AKT by 30%. We studied the inhibition of drugs using organoids of 1138 cells of LNCaP prostate cell line for 7 days with six drugs (see Table 1) and without any drug in a three-dimensional space with PhysiBoSS 2.0. We tested five different drug concentrations for each drug corresponding to the IC10, IC30, IC50, IC70 and IC90. In all the *in-silico* experiments, drugs diffused uniformly from the simulation boundaries.

Finally, in order to validate simulation results, we took a snapshot of the current state of the simulation every 240 min to create a growth curve for each simulation (Supplementary Figure S5). The median area under the curve (AUC) for 10 replicates was used to compare growth behaviours upon drug administration. Based on these AUC we defined a Growth Index as follows:

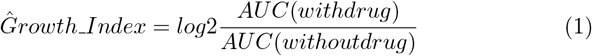

A Growth Index below zero indicates a reduction in growth upon drug treatment with respect to the untreated condition while a value over zero indicates an increase in growth.

### Double drugs studies

#### Estimation of drug-response for single and double drug studies

The effect of the drug on the cell depends on the concentration and provides insight into multiple drug characteristics such as potency or efficacy [32]. Usually, this relation exhibits a non-linear relation, and thus, sigmoidal-shaped functions such as the Hill equation are used to model the dose-response. Herein, we have used standard pharmacodynamics methods to model the effect of various concentrations of drugs on cell line growth. For this purpose, was used GDSC dose-response data (see Table 1) to fit the raw cell viability experiments using a multi-level fixed effect model [33] implemented in the *gdscIC50 R* package (https://github.com/CancerRxGene/gdscIC50). As a result, we obtained a specific sigmoidal dose-response curve for each drug and cell line pair (see Supplementary Figure 4).

Drug synergies were studied using the Bliss independence model [34] which is based on the idea that the two studied compounds are acting independently from each other, meaning that they are non-interacting [35]. Based on the effects of every single drug, a reference model was calculated as follows:

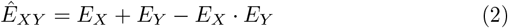

where *Ê*_*XY*_ is the predicted combined effect of how the two drugs *X* and *Y* act if no synergy or antagonism would exist; whereas 0 ≤ *E*_*X*_, *E*_*Y*_ ≤ 1 are the single drug effects of *X* and *Y*, respectively. If the measured combined drug effect observed is higher than the predicted effect *Ê*_*XY*_, synergy is declared, and antagonism is concluded otherwise. This can also be expressed using the Combination Index (CI) [36] which is calculated as follows:

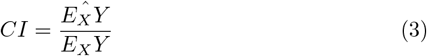

Where *E*_*X*_ and *E*_*Y*_ are the efficiency of the single drug inhibitions and *E*_*X*_*Y* is the inhibition resulting from the double drug simulations. A Combination Index (CI) below 1 indicates synergy while a value above 1 indicates antagonism.

#### Computational resources

All the simulations done in this paper were performed in the MareNostrum 4 supercomputer, located at the Barcelona Supercomputing Center in Spain. Each node contains two Intel Xeon Platinum 8160, each one with 24 processors running at 2.1 GHz and 33 MB L3 Cache. Memory is organised in two NUMA sockets with a total amount of 96GB per node. Individual simulations were done using all the 48 CPUs of one compute node. Model exploration performed using EMEWS was carried out using 10 nodes allocating three instances per node and assigning 16 CPUs per simulation instance. Results were analysed using custom scripts written in Python (3.8) language.

The tools and their repositories used in the present work are depicted in Table 2.

**Table 2.** Codes considered in present work with their maintainers and repositories

| Name | Repository | Maintainer |
| --- | --- | --- |
| PhysiCell | https://github.com/MathCancer/PhysiCell | Indiana University |
| MaBoSS | https://github.com/maboss-bkmc/MaBoSS-env-2.0 | Institut Curie |
| PhysiBoSS 1.0 | https://github.com/PhysiBoSS/PhysiBoSSv1 | Deprecated |
| PhysiBoSS 2.0 | https://github.com/PhysiBoSS/PhysiBoSS | BSC, Institut Curie |
| PCTK | https://github.com/PhysiBoSS/pctk | BSC |

## Results

In this section, we first introduce the redesign of PhysiBoSS together with the newly implemented features and functionalities. Secondly, we present novel results as examples of the kind of experiments possible with PhysiBoSS 2.0. We showcase here two examples: the integration of different cell receptor models for coupling the presence of environmental signalling substrates to the intracellular Boolean model as well as the integration of pharmacodynamics and Boolean models to conduct *in-silico* drug screen studies.

### The new design of PhysiBoSS 2.0 allows for extended functionalities

PhysiBoSS is a multiscale multicellular simulation framework that integrates PhysiCell [15] and MaBoSS [19, 20], enabling the simulation of cell signalling and gene regulatory networks in each individual cell agent. In this way, cell agents can integrate environmental and genetic signals and respond according to their internal Boolean model dynamics (see Figure 1A). Therefore, PhysiBoSS bridges the molecular level description of cell signalling and gene regulation with the surrounding microenvironment and the population dynamics, allowing us to explore the coupling between these different scales (see Figure 1B). For instance, PhysiBoSS 2.0 enables simulating drug studies such as adaptive therapy in cell populations while considering the role of heterogeneity among the cells and the environment and their role in the emergence of drug resistance [23].

With this new implementation, we aim to overcome several design problems from the original PhysiBoSS 1.0 [18]. In its original version, PhysiBoSS was implemented merging MaBoSS and PhysiCell by adapting several core classes of the latter, resulting in a new standalone simulation framework. This design proved to be very difficult to maintain and to keep aligned with the latest version of the original software components. For the same reason, bug fixing and the development of new features was also a difficult task. To solve these issues, PhysiBoSS 2.0 was redesigned and re-implemented from scratch as an add-on interface which decouples PhysiCell and MaBoSS minimizing dependencies and fixing many design problems of version 1.0 (Figure 1A and Supplementary Figure S1). The new add-on-based design simplifies the tool’s maintenance and allows for the independent upgrade of the different components, something not possible in the previous work of PhysiBoSS (see Supplementary Figure S2).

PhysiBoSS 2.0 uses the latest versions of MaBoSS and PhysiCell and thus incorporates all the new features and functionalities provided by the individual frameworks. For instance, by using PhysiCell 1.9, it is possible to track substrates internalised within cells and use these quantities as signals or inputs of a Boolean model; furthermore, it is also possible to implement rule-based cell-cell and cell-extracellular-matrix contact behaviours that trigger signals that propagate through a Boolean model [37]. On the other hand, the latest MaBoSS release (2.4.0) allows working with Boolean models of unlimited size as well as having these models defined in the SBML-qual standard (Supplementary Information). Among other new features, it is now possible to set up simulations (in the XML configuration file, see options in Table 3) which include different cell types that are associated with specific Boolean models. Moreover, we have standardised the integration of the Boolean model by using custom modules that connect intracellular variables to agent-based ones (discussed in section 5). Table 3 summarise, all the new features that can be defined by the user using the extended PhysiCell XML configuration file.

**Table 3.**
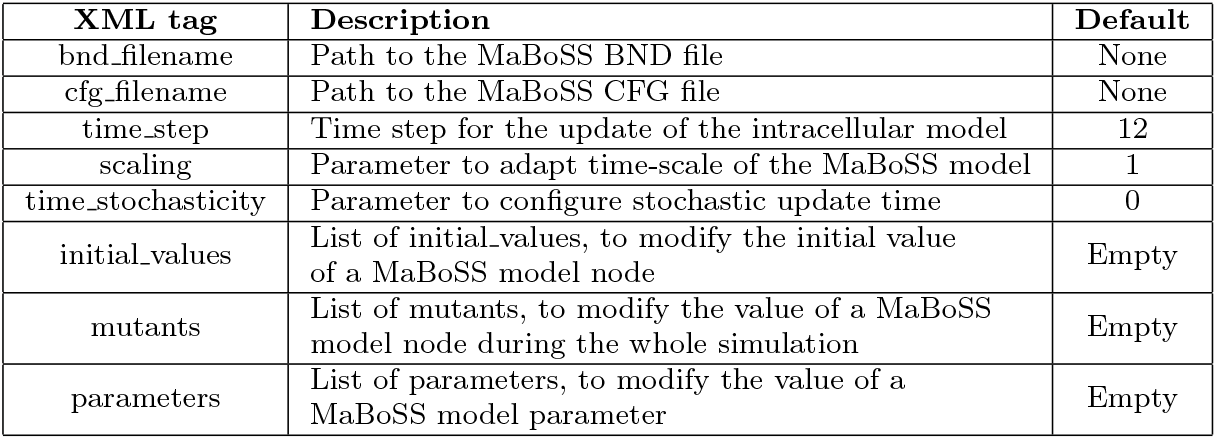
PhysiBoSS intracellular model configuration

| XML tag | Description | Default |
| --- | --- | --- |
| bnd_filename | Path to the MaBoSS BND file | None |
| cfg_filename | Path to the MaBoSS CFG file | None |
| time_step | Time step for the update of the intracellular model | 12 |
| scaling | Parameter to adapt time-scale of the MaBoSS model | 1 |
| time_stochasticity | Parameter to configure stochastic update time | 0 |
| initial_values | List of initial_values, to modify the initial value of a MaBoSS model node | Empty |
| mutants | List of mutants, to modify the value of a MaBoSS model node during the whole simulation | Empty |
| parameters | List of parameters, to modify the value of a MaBoSS model parameter | Empty |

Finally, different sample projects and templates are provided with the source code to facilitate the implementation of new models (see Section S2.2 in the Supplementary Text). PhysiBoSS 2.0 was written in C++ and the implementation is described in further detail in the Supplementary Information, together with the introduction of a template project for new users. PhysiBoSS 2.0 code is open-source distributed under BSD 3-clause license (https://github.com/PhysiBoSS/PhysiBoSS). PhysiBoSS 1.0 is also available at the following link: https://github.com/PhysiBoSS/PhysiBoSSv1. In addition, we also provide a nanoHUB GUI-based tool implementing an example model, to ease its use at https://nanohub.org/tools/pba4tnf/.

#### Handling and processing simulation outputs

Processing, analysing and visualising multi-scale simulations’ output is a non-trivial task; it requires handling and integrating several output file formats into aggregated easy-to-use data frames as well to use 3D rendering libraries to generate visual representations of the system. For this reason, together with PhysiBoSS 2.0, we have developed the PhysiCell ToolKit (PCTK) a python-based package that includes a library and command-line scripts to process and analyse simulation outputs. Although there are already available tools for handling PhysiCell outputs (such as https://github.com/PhysiCell-Tools/python-loader), with PCTK we aim to gather and organise different pieces of python code that have been recurrently useful in different projects involving PhysiCell and PhysiBoSS. Currently, the package implements a module to parse and handle the MultiCellDS standard and uses an efficient schema to process the output files containing the cells and microenvironment data. Moreover, on the top of this module, we have implemented different command-line tools for processing and creating basic plain-text, data-frames and generic plots, including time courses of the number of alive and dead cells; it also provides a command-line tool to allow users to generate POV files used as inputs for the 3D rendering of the multicellular models using POV-Ray (Persistence of Vision Raytracer, PovRay 2004). PCTK can be used both as a callable library as well as a stand-alone command-line tool and can be found at https://github.com/PhysiBoSS/pctk. Documentation and examples are also provided on the PCTK site.

### PhysiBoSS 2.0 reproduces original results and allows to test new transport mechanisms

The new functionalities of PhysiBoSS 2.0 allow for better modularity and ease of reuse of functions. These allow, for instance, researchers to focus on the important step of defining the interactions between variables of the intracellular Boolean models and of the cell agent and environment part.

Depending on the level of detail desired different approaches can be used, but in general, the problem can be split into two different sub-problems: how to couple cell and environmental signals (continuous variables) into the Boolean input nodes, and how to use the value of the Boolean output nodes to control the behaviour of the cell agents. In general, the mapping of continuous variables into a Boolean value requires the use of a transfer function. The simplest case is the use of a step function *H*(*x*) which returns 1 when *x* ≥ *θ* and 0 otherwise, for a given threshold *θ*, as we had in [18]. On the other hand, using the value of the Boolean model’s output node to control the cell agent behaviour can be done by modulating the specific rates of the cell cycle or death models, as well as by triggering custom rules defined by the modeller (see for example [37]).

In order to test and validate the new PhysiBoSS 2.0 and to show how the different submodels can be integrated, we have re-implemented the different models presented together with the original PhysiBoSS 1.0 and replicated all the results reported by [18]. Letort et al. (2019) implemented a multi-scale model of 3T3 fibroblast spheroids to investigate the complex dynamics observed when tumour cells are exposed to different regimes of tumour necrosis factor (TNF) [38]. The model integrates the Cell Fate Boolean network [14] inside the cell agents to simulate the growth of a spheroid of cancer cells under different treatment regimes, which correspond to the supply of TNF pulses with different frequencies, duration and concentrations of the cytokine.

We re-implemented the model in PhysiBoSS 2.0 using an extended version which includes a more detailed description of the TNF-receptor binding mechanisms [23]. Figure 2 shows a schematic representation of the multi-scale model as well the difference between the TNF receptors, and growth model used in [18] and [23]. Details of the model formulation and implementation can be found in the supplementary Text. The experiments consist of an initial tumour spheroid of 1000 cells that are exposed to different concentrations and regimes of TNF. We run all the different experiments using the same simulation setups to replicate Figure 4 from [18] using both,the re-implemented model and PhysiBoSS 2.0 and the original code. The comparative results are presented in Figure 2, where each panel shows the output generated by each of the two PhysiBoSS versions.

**Fig. 2.**
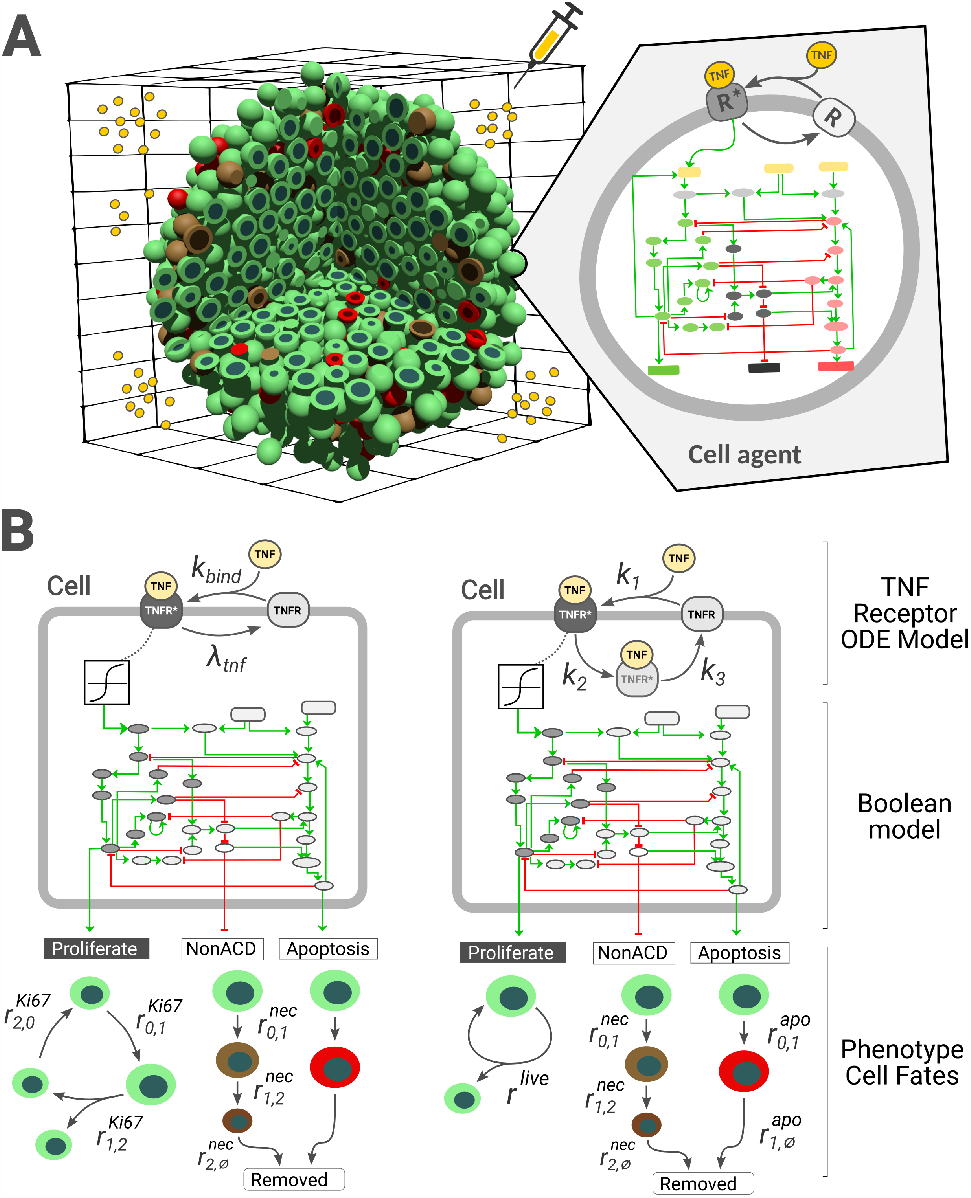
Different implementations of the multi-scale TNF models. Panel A schematically represents the multi-scale model used to explore the 3T3 fibroblast spheroids growth dynamics under different TNF exposure regimes. Panel B shows the intracellular models used in the PhysiBoSS original publication [18] (left) and the extender version implemented in PhysiBoSS 2.0 (right).

The results show that the simulations obtained with PhysiBoSS 2.0 qualitatively reproduce the results reported by Letort et. al (2019). Specifically, the new implementation correctly reproduces the pattern observed when cells are exposed to a continuous supply of TNF causing cells to become resistant to the effect of the cytokines (Figure 3B-D). Moreover, the model implemented in PhysiBoSS 2.0 also correctly predicts the reduction of the tumour when cells are exposed to short pulses of TNF at a frequency of 150 minutes (Figure 3E) as well as the infective regime of short pulses of TNF at a frequency of 600 minutes (Figure 3E). Furthermore, we can observe that PhysiBoSS 2.0 better captures the exponential cell growth that was somehow biphasic in PhysiBoSS (Figure 3A). The differences observed in all the experiments are mainly due to the differences in how the TNF binding and the cell cycle models were implemented in the new version (see Figure 2B). Beyond the difference in the transport mechanism, the new version was implemented with a clear separation of the different submodels as well as the interfaces to connect them, and thus it can be used as a sample project to implement new models since the code is packed with PhysiBoSS 2.0.

**Fig. 3.**
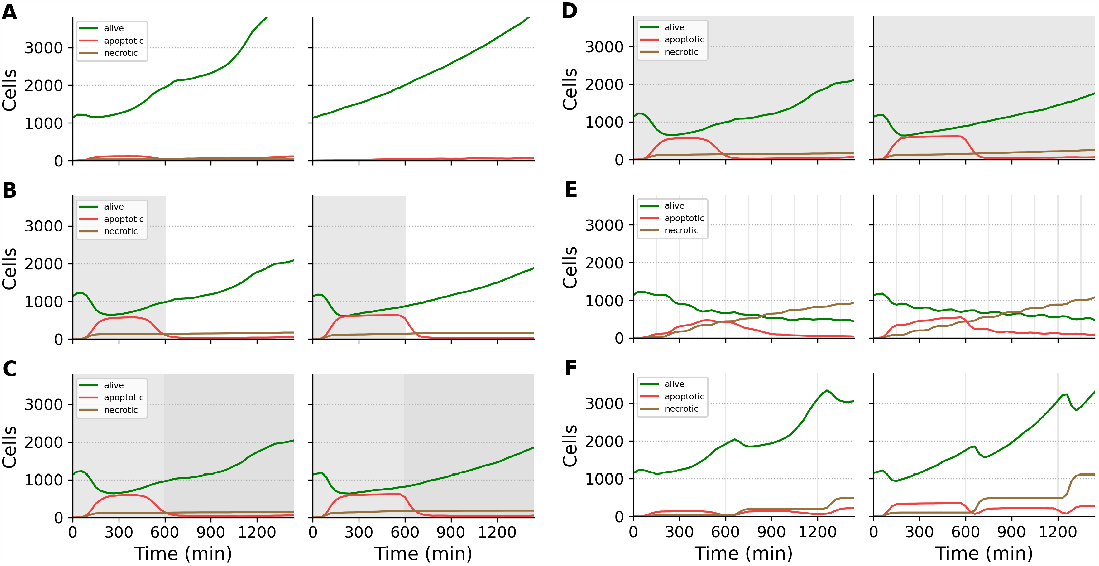
Pair comparison between results obtained using the original PhysiBoSS (left side) and PhysiBoSS 2.0 (right side). The plots represent population growth curves for the same TNF pulse in-silico experiments reported in the original PhysiBoSS publications. Each panel corresponds to a different *in-silico*experiment. A) no TNF added; B) single pulse of 0.5 ng/mL for 10 hours (600 min); C) single pulse of 0.5 mg/mL for 10 hours followed by a second pulse of 5 mg/mL for 14 hours; D) continuous pulse of 0.5 ng/mL throughout the 24h the experiment last (1440 min); E) TNF pulses of 0.5 ng/mL and duration of ten minutes at intervals of 150 minutes; and F) TNF pulses of 0.5 ng/mL and duration of ten minutes at intervals of 600 minutes. Vertical grey patches represent the TNF pulses.

### Simulating drug screening studies and drug synergies

We used PhysiBoSS 2.0 new functionalities to portray the typical phenotypic characteristics of cell lines and its drugs’ effects and synergies by using personalised Boolean models [24, 28] and GDSC dose-response profiling data [33]. The models described in the previous sections rely on a dynamic model that mechanistically describes the interaction between the signal molecule (TNF) and its target (TNF-receptor) and a step function to transfer the level of TNF bound in the membrane to the corresponding Boolean input node. However, the detailed mechanism and kinetics of how drugs interact with their targets are largely unknown. For such cases, in PhysiBoSS 2.0 users can leverage experimental dose-response curves, such as the ones from GDSC, to include the effect of drug concentrations to inhibit a node of the intracellular signalling model.

Therefore, when a drug is available at a specific concentration in the surroundings of a cell, the cell response (its viability in this case) is looked up in the dose-response curve that is specific to the cell line and the drug. Then, the corresponding drug target node in the Boolean model is inhibited with a probability of 1−*cell viability* (see Material and Methods for further details). This ensures that for a concentration of IC90, for example, a node inhibition would be obtained in 90% of the cases and that an IC10 would lead to a node inhibition with just 10% probability (see an example dose-response curve in Supplementary Figure S4). Here, we showcase examples that integrate pharmacodynamics and Boolean models to study single drug effects, drug combinations and their synergies.

#### Single drug screening studies

Using PhysiBoSS 2.0, we simulated organoids of 1000 cells of the prostate cell line LNCaP for seven days with six drugs (Table 1) in a three-dimensional domain where the drugs were supplied from the domain boundaries (the box walls) and diffusing to the centre of the simulation space (Figure S12). We tested five different concentrations for each drug corresponding to the IC10, IC30, IC50, IC70 and IC90. This rather simple setup leads to varying drug availability and effects throughout the different layers of the tumour and, therefore, ends up affecting heterogeneously the different cancer cells depending on their location and if they are surrounded by cells or open to the microenvironment (see Section 5).

We simulated the five concentration simulations for all six drugs (see the growth indexes values in the diagonal of Figure S6) and performed a Kruskal-Wallis rank sum test to analyse the significance of growth behaviour changes for each drug simulation, finding two out of six drugs significant: Ipatasertib (targeting AKT) and Pictilisib (targeting PI3K) with *P* ≤ 0.0001 (Table S2). The growth indices for these two significant drugs are shown in Figure 4B and C in the first column and row indicated by “None”. For instance, when simulating Pictilisib dosages, we can see that starting with IC10 there is a decay in the Growth Index that is maximal when reaching IC70 and IC90 Figure 4B.

**Fig. 4.**
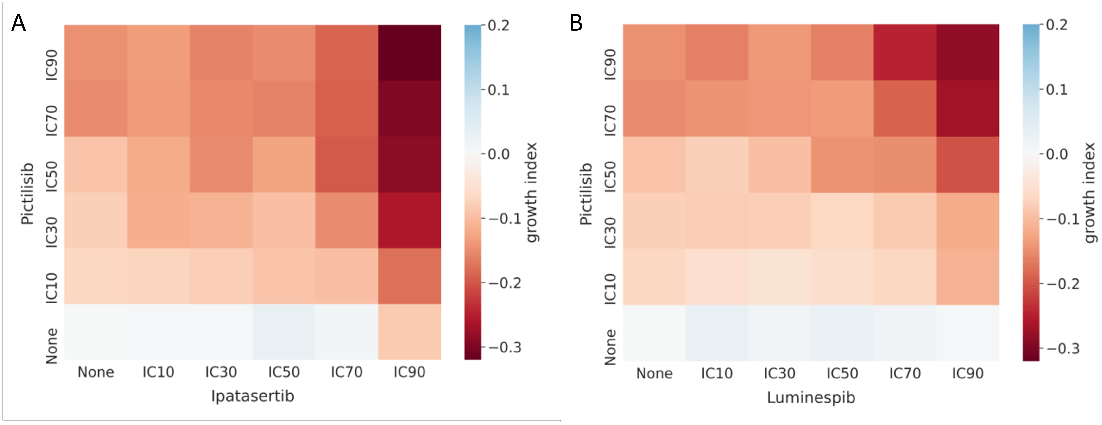
Growth index of the multiscale simulations with Pictilisib + Ipatasertib (A) and Pictilisib + Luminespib (B) with respect to the untreated LNCaP. Each simulation was replicated 10 times. For each combination, the growth index was obtained by taking the log2 of the ratio between the median AUC upon drug administration and the median AUC of the untreated simulations. White colour means no growth behaviour change upon drug administration, blue means the drug increased the growth and red means that the drug diminished the growth of the cells.

#### Double drugs screening studies

Furthermore, PhysiBoSS 2.0 also enables combining the administration of more than one drug allowing the simulation of drug-synergy studies. As with the single-cell studies, we used PhysiBoSS 2.0 to simulate spheroids of 1000 LNCaP cells with combinations of the aforementioned six drugs with their five different concentrations (Table 1). We identified two combinations of drugs as interesting: Pictilisib with Ipatasertib and Pictilisib with Luminespib (targeting HSPs). For both drug combinations we can observe a gradual growth inhibition with respect to the drug concentration and, furthermore, the indication of drug synergy as combined inhibition levels outdo single effects. While the drug combination Pictilisib and Luminespib requires high drug concentrations such as IC70 and IC90 for both drugs to obtain a Growth Index of around -0.3, the drug combination Pictilisib and Ipatasertib reaches similar values already with much lower concentrations for Pictilisib such as IC30 or IC50 (Figure 4B, C and Supplementary Figure S6).

In addition, we studied the Bliss independence of these combinations (Figure 5 and Supplementary Figure S8) and found complex synergies in both cases. Synergy values reach their maximum for Pictilisib and Luminespib with high drug concentration values of both compounds. Especially high drug concentrations of Luminespib drive the synergy value of the drug combination. For Pictilisib and Ipatasertib, on the other hand, synergy values peak for IC values around IC50 and IC70 for both compounds. Comparably, high drug concentrations of Ipatasertib especially drive the synergy values. For both drug combinations synergy values strongly decrease or even turn into slight antagonism with low drug concentrations for all drugs. Such varying combinatorial effects depending on the drug concentration have previously been observed. Drug combinations can be both antagonistic and synergistic at the same time which can be described with the help of exposure-response surfaces [39]. These surfaces can help in choosing drug concentrations for combinatorial therapies. The complete study of LNCaP and six drugs (Ipatasertib targeting AKT, Luminespib targeting HSPs, Pictilisib targeting PI3K, Afatinib targeting EGFR, Ulixertinib targeting MAPK1, Selumetinib targeting MAP2K1 and MAP2K2) can be found in the Supplementary Material, Section S4.6.

**Fig. 5.**
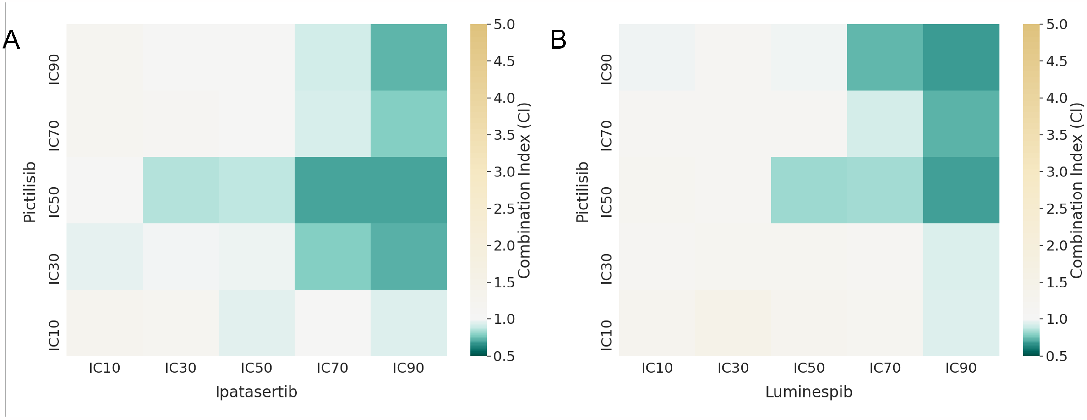
Bliss independence Combination Index (CI) of the multiscale simulations of LNCaP with Pictilisib and Ipatasertib (A) and Luminespib and Pictilisib (B). White colour indicates an additive effect, green colour a synergistic effect, and yellow an antagonistic effect. For a complete figure of all the combinations refer to S8.

#### Studying the heterogeneity of drug screening simulations

The integration of drug simulations in PhysiBoSS allow to expand the study of the heterogeneous behaviours emerging from the cells. In Letort et al. (2019) [18], we studied the effect of different genetic backgrounds, such as mutants cFLIP+ IKK+ and CASP3+ Cytc+, in their response to TNF treatments, as we have replicated in PhysiBoSS 2.0 (Sections S4.8 and S5.3 and Figure S14). In addition to the genetic perturbations, PhysiBoSS 2.0 can be used to study the heterogeneity in the cells’ response to a drug inhibition in a population of cells that share a common genetic background. In this scenario, heterogeneity comes from a non-homogeneous microenvironment and is caused by the drug penetration into the cell population resulting in varying substrate availability in different regions of the microenvironment that affect the drug’s effectiveness.

In our simulations, the drug is supplied on the simulation boundaries assuring a uniform administration of the drug. As cells act as sinks for the drug, by uptaking it at a dynamic rate, cells in the outskirts of the spheroid will have more available drugs than the ones in the spheroid centre (see Supplementary Figure S12).

To study the effect of the position of the cell on their response to the drugs, we simulated a spheroid of a radius of 100 *μm* with drug administration on the boundaries with 10 replicates. We studied the growth rate of the different layers of the spheroid by splitting the spheroid into 50 *μm*-thick layers from their Euclidean distance to the tumour centre. At the beginning of the simulations, the cells are only in the inner layers 1 and 2 as the initial radius is 100 *μm*. Then the cells start expanding into layer 3 at around 200 minutes in the simulation, then layer 4 at around 5000 minutes and then 5 at around 8000 minutes.(Figure 6A) depicts the number of cells for each spheroid layer of one simulation with and without drugs, but these were consistent across the replicates.

**Fig. 6.**
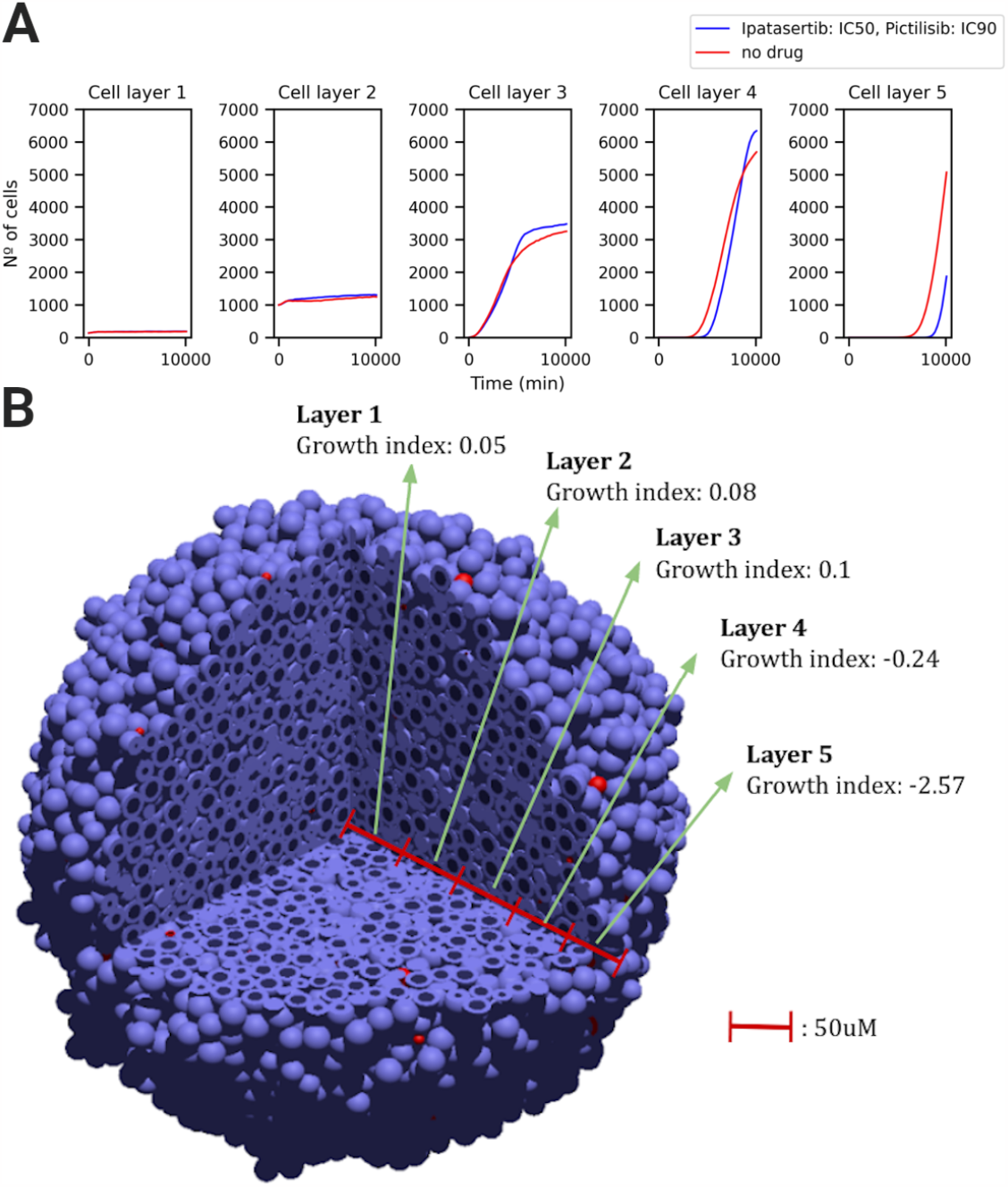
Studying the heterogeneity of drug screening simulations. Panel A shows the growth curves for the no-drug simulation and the drug simulation with Ipatasertib(IC50) + Pictilisib (IC90) separated into sphere layers. By taking the distance to the tumour centre we define 5 50 *μm*-thick layers: layer 1 is the innermost layer and layer 5 corresponds to a distance from 200 to 250 *μm* from the tumour centre. From these growth curves the AUC values and growth indices are calculated for each layer separately. Panel B shows a 3D representation of the cell population growth indices of the 5 50 *μm*-thick layers at the end of the simulation. Layer 1 is the innermost layer from the centre of the tumour to a distance of 50 *μm* to it. Blue cells are living cells; Red cells are apoptotic.

The growth curves obtained when applying Ipatasertib (IC50) + Pictilisib (IC90) (Figure 6A, blue) and when applying no drug (Figure 6A, red) are almost identical for the inner layers 1 and 2, indicating the absence of drugs in the central tumour regions. Drugs have a deleterious effect starting at layer 4, where we see that the cells from the no-drug simulation reach before the beginning of this layer (at 150*μm* from the centre) causing the red no-drug curve to start before the blue one. This difference increases in layer 5, where the red no-drug curve starts much sooner than the blue drug curve. In these layers, the cells are closer to the drug source and are uptaking more drug that is killing them by apoptosis.

Interestingly, the reduced survival of layers 4 and 5 of the drug simulations is causing a slight increase in growth rate in layers 1, 2 and 3 when compared to the no-drug simulation. We think this increased growth in the inner layers can be explained by the increase in available space. As drugs reduce the number of cells in outer tumour regions, it leaves more space for cells in inner tumour regions to have space to proliferate. In fact, we can observe in sphere layers 3 and 4 the intercept point of when the tumour reaches a size when not enough drug can reach the cells and the reduced amount of surrounding cells increases the available space to divide. finally, the no-drug simulations have scarce space in the inner layers, causing the cells to have an outwards pushing dynamic that makes them reach layers 4 and 5 much sooner than the drug simulations. We gathered the layer-specific Growth Index by calculating the AUCs of the median of the 10 replicates and saw that an administration of drugs on the boundaries leads to varying growth indices among the tumour sphere layers (Figure 6B). Cells located in the central tumour regions do not show any signs of growth inhibition as they have slightly positive growth indices whereas cells on the tumour boundaries show high growth inhibition values with negative growth indices.

In summary, the results of these experiments respond to two different dynamics: first, by uptaking drug from the environment, cells from the outer layers have a shielding or protective effect on cells from the inner layers; second, the death of the cells exposed to drugs in the outer layers free up space for the proliferation of cells from the inner layers, even in the innermost layer where space is the scarcest.

## Discussion

The new add-on design of PhysiBoSS 2.0 allows us to extend its functionalities and reuse in a more efficient way compartmentalised functions and replicate simulations [25]. This modular design ensures long-term sustainability as either tool (PhysiCell and MaBoSS) can now be updated independently (Figure S1). Furthermore, we also include several sample project with different degrees of complexity than can be used as templates to develop novel models. In addition, we also provide the PCTK python package to ease the task of handling and processing simulation outputs and to easily create summary plots and 3D rendered figures.

As a showcase example, we have here presented a drug screening study that uncovered synergies between drugs and heterogeneities in the cell population and that expands our work done on prostate Boolean models [24]. We have presented drug screening studies that could mimic, albeit simplistically, *in-vitro* spheroid experiments and have validated some of them using real-time cell survival assay data. Implicitly, this work also showcases the flexibility of PhysiBoSS 2.0 to run with any user-provided Boolean model in MaBoSS or SBML-qual format, as opposed to the original PhysiBoSS.

As drug administrations are never completely homogeneous but rather occur from the tumour surroundings or from specific points in the tumour environment, such as blood vessels, we consider it of high importance to continue studying these growth heterogeneities and the shielding effect that cells from the outer layers can have on other cells from the inner layers to deliver a realistic digital twin of drug treatment.

Additionally, we present ways of simulating two potential causes of tumoural heterogeneity: the uneven drug penetration in the tumour caused by the microenvironment and the different genetic backgrounds of the tumour cells. The study of this kind of heterogeneities is needed to have real-size digital twins [26] that allow for the understanding of the mechanisms behind drug treatment evasion. Current approaches to predict drug synergies do not consider the complexity and heterogeneity typically found in a tumour, even though it is known that these greatly affect the drug effectiveness [40–42].

With these added capabilities, PhysiBoSS 2.0 takes important steps towards having truly personalised digital twins that could be used for realistic clinical trials of drug treatments and effective drug synergies. We are currently working on expanding this tool to include the extracellular matrix and its interactions with the cells [37], blood vessels and their vascularisation and complex three-dimensional architectures taken from spatial omics of patients.

## Conclusion

PhysiBoSS 2.0 is a redesign and reimplementation of PhysiBoSS to be a reusable, extensible and updatable add-on for the PhysiCell framework. Our tool follows a modular design that decouples PhysiCell and MaBoSS codes, minimising the dependencies between these tools and ensuring long-term maintainability and is, therefore, more respectful of the single responsibility principle and the façade pattern software design [43].

We show added functionalities that enable PhysiBoSS 2.0 to study disease mechanisms that cause malfunctions in personalised models while being flexible to handle many different models and consistent with the results of the former PhysiBoSS. Altogether, the new design and implementation allow PhysiBoSS 2.0 to be model-agnostic, and easily customisable by users and provides a simple framework of custom modules and custom settings to use with any Boolean model of interest in MaBoSS or SBML-qual format. We have made efforts to provide full accessibility to PhysiBoSS 2.0 code as well as to several accompanying interoperable tools that make the full software bundle much more reusable at https://github.com/PhysiBoSS/.

## Supplementary information

If your article has accompanying supplementary file/s please state so here.

Authors reporting data from electrophoretic gels and blots should supply the full unprocessed scans for key as part of their Supplementary information. This may be requested by the editorial team/s if it is missing.

Please refer to Journal-level guidance for any specific requirements.

## Acknowledgments

Acknowledgments are not compulsory. Where included they should be brief. Grant or contribution numbers may be acknowledged.

Please refer to Journal-level guidance for any specific requirements.

## Declarations

Some journals require declarations to be submitted in a standardised format. Please check the Instructions for Authors of the journal to which you are submitting to see if you need to complete this section. If yes, your manuscript must contain the following sections under the heading ‘Declarations’:

- Funding
- Conflict of interest/Competing interests (check journal-specific guidelines for which heading to use)
- Ethics approval
- Consent to participate
- Consent for publication
- Availability of data and materials
- Code availability
- Authors’ contributions

If any of the sections are not relevant to your manuscript, please include the heading and write ‘Not applicable’ for that section.

Editorial Policies for

Springer journals and proceedings: https://www.springer.com/gp/editorial-policies

Nature Portfolio journals

https://www.nature.com/nature-research/editorial-policies

*Scientific Reports*

https://www.nature.com/srep/journal-policies/editorial-policies

BMC journals

https://www.biomedcentral.com/getpublished/editorial-policies

## Section title of first appendix

An appendix contains supplementary information that is not an essential part of the text itself but which may be helpful in providing a more comprehensive understanding of the research problem or it is information that is too cumbersome to be included in the body of the paper.

