## Supplementary File 1 for "PhysiBoSS 2.0: a sustainable integration of stochastic Boolean and agent-based modelling frameworks"

### S1 Extended Materials and Methods

#### S1.1 MaBoSS

MaBoSS [1, 2] is an open-source standalone software package, as well as a library written in C++, for simulating Boolean models using a continuous/discrete-time Markov processes approach. A Boolean network can either represent cell signalling pathways or any regulatory network expressed as a Boolean model [3]. MaBoSS uses transition rates to each node transition and, given some initial conditions, MaBoSS applies Monte-Carlo kinetic algorithm [4] to the Boolean transition state space to evaluate temporal trajectories. Some examples of the use of stochastic Boolean simulations are uncovering transient states between invasive and migration phenotypes in breast cancer [5] and examining combinations of genetic alterations in bladder cancer [6].

For this project, we are using MaBoSS 2.4.0 and its source code is available at: https://github.com/maboss-bkmc/MaBoSS-env-2.0

##### S1.2 PhysiCell

PhysiCell [7] is an open-source multiscale multicellular simulation framework written in C++. It was developed as an agent-based modeller where the agents are cells with properties like secretion/uptake values for different substrates, mechanical properties, growth rates, who interact with the tissue environment and with the other cells. PhysiCell runs over BioFVM [8], providing the basic structure for the agents, and performing the calculus for diffusion, update and secretion of different molecules available in in the environment. Some examples of the use of PhysiCell are discovering inmune-tumour interactions [? ] or target delivery of therapeutic compounds to tumour cells [9]. For this project, we are using PhysiCell version 1.9.0 and its source code is available at: https://github.com/MathCancer/PhysiCell

##### S1.3 PhysiBoSS 1.0

PhysiBoSS 1.0 [10] is an open-source multiscale multicellular standalone simulator created by using PhysiCell and MaBoSS. PhysiBoSS added an additional time scale to the default PhysiCell, adding cell signalling to each agent. This allows the agents to integrate environmental and genetic signals and respond according to their Boolean model. PhysiBoSS was used to study the heterogeneity of a cell population in response to different TNF treatments and their population dynamics. This software is available at: https://github.com/gletort/PhysiBoSS.

#### S2 PhysiBoSS 2.0 implementation as an add-on of PhysiCell

The original PhysiBoSS [10] interacts with MaBoSS using mainly two classes: i) the MaBoSSNetwork class; and ii) the `CellCycleNetwork` class. The `MaBoSSNetwork` class is the interface that communicates with the MaBoSS engine and the `CellCycleNetwork` is the class in charge of preparing and performing the MaBoSS simulations and interacting with PhysiCell variables according to the resulting cell fate after the MaBoSS simulation. In PhysiBoSS, these two classes are tightly connected with PhysiCell core classes. However, we reused these classes' structure and a lot of their functionalities for our add-on. Next, we refactored the classes of our add-on. The `MaBoSSNetwork` class required a minor refactoring: trimming the specific functions used in PhysiBoSS and updating the MaBoSS simulation engine to allow for simulations to be run in parallel.

On the other hand, `CellCycleNetwork` required much more work as it is the interface with PhysiCell. For this part, we wanted to help develop an interface for intracellular models into the main PhysiCell code and provide

MaBoSS intracellular models as a first example. After discussions with PhysiCell's developers, we decided to represent intracellular models as part of the PhysiCell's Phenotype class, which is a component of all cells. This allows us to have our models associated with cell types and defined in the newly introduced cell definitions in the PhysiCell XML settings.

The decoupling is obtained by encapsulating the functionalities used to simulate the Boolean model in a dedicated `MaBoSSNetwork` class that uses the MaBoSS library to store the current Boolean state of the cell and perform the simulations. This class is then used in a novel of PhysiCell's Intracellular interface (`MaBoSSIntracellular`), which implements the connection to the MaBoSS intracellular model that provides the functionalities to simulate the Boolean model.

We started by creating a pure abstract class called `Intracellular`, defining a simple interface to control our intracellular submodels. We then created a new `MaBoSSIntracellular` class in our PhysiBoSS folder, implementing the methods of the Intracellular interface. Such methods include loading model description in the XML settings, initialising the model, updating the model or getting/setting values of the variables of the model. Some functions from the `CellCycleNetwork` class of PhysiBoSS 1.0 were reused; nonetheless, any control over the life and death models were remove from PhysiBoSS 2.0 core. Moreover, this also allow to let the definition of the interaction between the different models to be specified at the level of each model.

#### S2.1 Design and implementation of a MaBoSS add-on for PhysiCell

We started the project by creating a fork of PhysiCell's [7] master branch from its repository (https://github.com/MathCancer/PhysiCell) to ensure we are using the latest release. Once we had our own copy of PhysiCell, we created a folder in the root of the project named add-ons, where all custom extensions of PhysiCell should be placed (see Figure S2). Figure S2 shows a schematic representation of how the project is structured; inside the add-ons folder, we set the folder PhysiBoSS, where all the code and dependencies necessary to run our add-on are located. On the one hand, the *src* folder stores the PhysiBoSS code that interfaces between PhysiCell and MaBoSS. Furthermore, when a PhysiBoSS project is instantiated, the latest MaBoSS release code is retrieved from its repository (currently: https://github.com/maboss-bkmc/MaBoSS-env-2.0) and compiled to be linked to PhysiBoSS projects.

Figure **??**B depicts a simplified version of the Unified Modeling Language (UML) class diagram representing the relationship between PhysiCell Intracellular interface, PhysiBoSS add-on and MaBoSS. This Intracellular interface has since been included into version 1.9.0 of PhysiCell and is already used by two other add-ons (https://github.com/MathCancer/PhysiCell/releases/tag/1.9.0 and its currently working with the last PhysiCell Release 1.10.4. This add-on design allows PhysiBoSS code to uncouple PhysiCell and MaboSS making

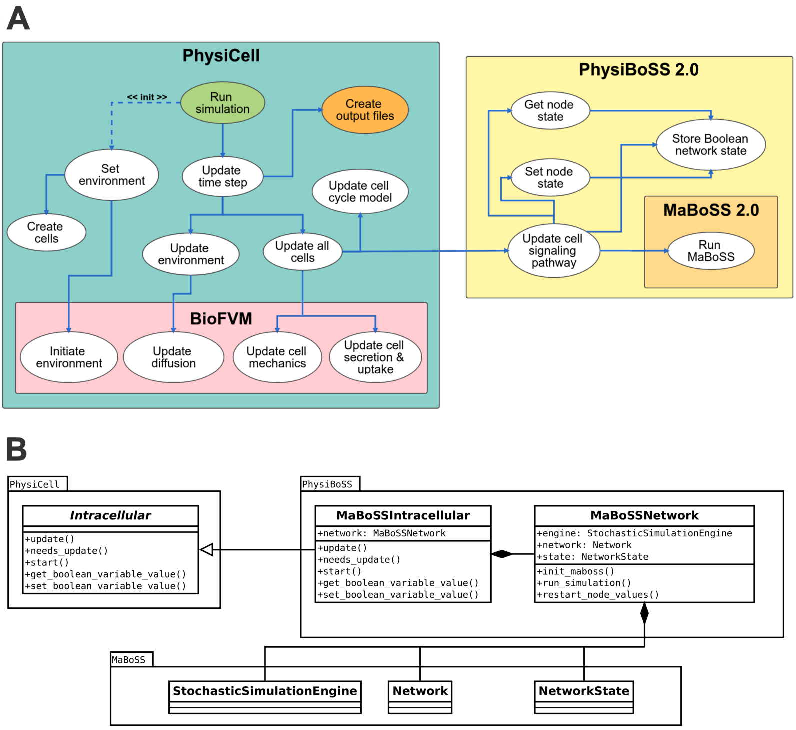


**Supplementary Fig S1 UML representation of PhysiBoSS 2.0 design.** Panel A shows a high-level view of the PhysiCell + PhysiBoSS 2.0, and the communication between the different components. Panel B shows the UML class diagram showing the communication between the modelling packages PhysiCell and MaBoSS through the interface defined by PhysiBoSS 2.0.

the whole project much easier to maintain; whenever new versions of PhysiCell or MaBoSS are included in PhysiBoSS, users may only have to adapt the MaBoSSIntracellular or MaBoSSNetwork class, respectively.

#### S2.2 Defining a template model connecting a Boolean model with PhysiCell

To facilitate the use and further development of PhysiBoSS 2.0, we have designed a template that connects a drug-responsive Boolean toy model with PhysiCell's live cell cycle. This template can be used as a basis to study how to connect the environment to the Boolean model and this to the agent behaviour:

https://github.com/PhysiBoSS/PhysiBoSS/tree/master/sample_projects_intracellular/boolean/template_BM.

```
/PhysiCell
├── addons
│   └── PhysiBoSS
│       └── src
│           ├── maboss_intracellular.cpp
│           ├── maboss_intracellular.h
│           ├── maboss_network.cpp
│           ├── maboss_network.h
│           ├── utils.cpp
│           └── utils.h
├── beta
├── BioFVM
├── config
├── core
├── custom_modules
├── ⋮
├── sample_projects_intracellular
│   └── boolean
│       ├── physiboss_cell_lines/
│       ├── prostate
│       ├── spheroid_tnf_model
│       └── template_BM
└── ⋮
```

**Supplementary Fig S2 PhysiBoSS project structure.** Simplified tree structure representation of PhysiCell and PhysiBoSS main folders and files.

Two drugs, Ainhib and Binhib are set in the file `PhysiCell_settings.xml` in `template_BM/config` folder (lines 217-230), they can be dosed in the environment constantly or with a given frequency (`PhysiCell_settings.xml`, lines 256-266), similarly to how it is done in the TNF model, described in the previous section. The values for the inhibitors are internalised (`template_BM/custom_modules/custom.cpp`, lines 147-164) and updated in each agent (`custom.cpp`, lines 77-90), and their internalised values determine the value of Boolean nodes *anti_A* and *anti_B* based on a user-defined threshold (`custom.cpp`, lines 153-163).

The nodes *anti_A* and *anti_B* inhibit nodes $A$ and $B$, respectively, inactivating $C$, which needs either $A$ or $B$ to be active (`template_BM/config/boolean_network/toyBM.bnd`). Whenever $C$ is active the cell division rate is multiplied by 20 (`custom.cpp`, lines 168-183). This cell division rate of the "live" cell cycle model can be further studied in Section 17.1.1 of PhysiCell's user guide in the documentation folder.

Altogether, modellers can easily define the rules that modify the state of the input nodes in the Boolean model based on inputs from PhysiCell, such as the internal concentration of different molecules, as well as other types of

**Table S1** Doubling times for the 6 prostate cell lines used with corresponding media composition and references.

| Prostate cell line | Doubling time (hours) | Media composition | COSMIC ID (GDSC) | Source |
|---|---|---|---|---|
| LNCaP | 42 | T medium, 5% serum | 907788 | link |
| 22Rv1 | 35 | T medium, 5% serum | 924100 | link |
| BPH1 | 50 | 80% RPMI 1640 + 20% h.i. FBS + 20ng/mL testosterone + 5 ug/m: transferrin + 5 ng/mL sodium selenite + 5 ug/mL insulin | 924105 | link |
| DU145 | 30 | DMEM, 10% serum | 905935 | link |
| PC3 | 30 | T medium, 5% serum | 905934 | link |
| VCaP | 53 | DMEM, 10% serum | 1299075 | link |

signals; and used the state of the Boolean model to trigger different behaviours of the cell agent, such start growing or entering into apoptosis.

### S3 Personalisation of cell lines in PhysiBoSS 2.0

Apart from using the personalisation of Boolean models using omics data from PROFILE [11] described in 5, we have also personalised the doubling times of the different prostate cell lines from literature experimentsS1.

### S4 Multiscale simulations of drug treatments and combinations

#### S4.1 Drug-target selection for LNCaP-specific Boolean models

In PhysiBoSS 2.0, diffusing drugs in the microenvironment target and inhibit a specific node in the Boolean model. These nodes represent proteins that are known to affect prostate cancer in this cell line, more information on the work from [12]. We used the Drugbank database [13] and matched with its availability on the GDSC database [14] to identify such target-node-drugs relationships of interest for LNCaP (Table 1).

Furthermore, we investigated the nodes that were inhibited by the drugs used in our study (Table 1). In a previous work [12], we studied the LNCaP Boolean model in detail with MaBoSS: we knocked out nodes by forcing their node value to 0 throughout the whole simulation. In the present work, this would correspond to a drug being fully effective, so being available to a cell at IC100 (see Supplementary Materials, Section S4.2). We then observed changes in phenotype probabilities for the readout nodes such as Apoptosis and Proliferation by comparing the knock-out model to the original LNCaP model. This phenotype change is expressed with the help of a phenotype variation index (Phenotype score of inhibited LNCaP - Phenotype score of wild-type LNCaP).

From our former results, we extracted the phenotype variations for the six single nodes corresponding to the drugs used in this work (Fig S3). We observe a similar pattern of phenotype variations for all six nodes, where all of them affect, at the same time, both Proliferation and Apoptosis. The node inhibitions that lead to the highest increase in apoptosis are AKT, ERK and PI3K. Furthermore, the ones that deplete proliferation the most are AKT, EGFR and PI3K. We can conclude that on the Boolean model level, the used drugs act through both the depletion of proliferation and the increase of apoptosis. In PhysiBoSS 2.0, the Boolean model outputs are mapped to the agent-based simulation affecting the growth rate of cells and cell death and, therefore, affect the overall cell number, the AUC and finally the growth index (see Section 5). Due to the fitting of these mapping functions, the proportions of proliferation depletion and apoptosis increase might differ between PhysiBoSS 2.0 and MaBoSS. Note that cells in our PhysiBoSS 2.0 setup rarely have access to a drug concentration corresponding to the IC100 attenuating the overall killing effect of the drug.

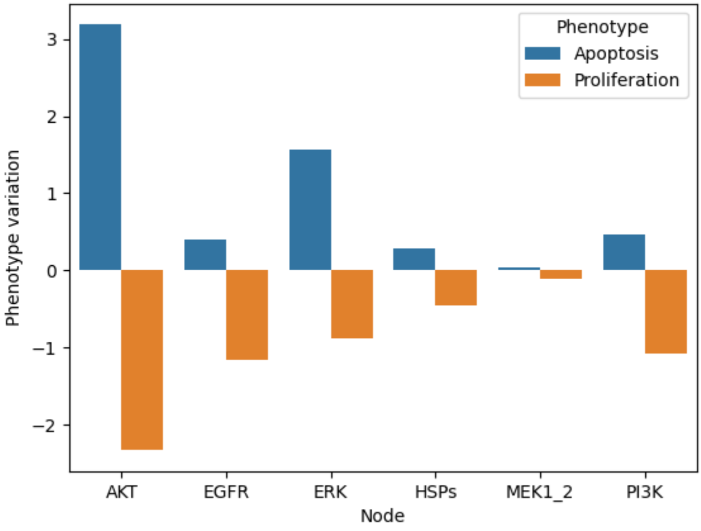


**Supplementary Fig S3 Apoptosis and Proliferation score variations upon single node inhibition (node forced to 0) for the 6 nodes corresponding to the drugs studied in this work, see Table 1.**

#### S4.2 Integration of dose-response curves

Experimental dose-response curves were fitted using the gdscIC50 package. The format for the integration of the dose-response curve into our framework is a CSV file that includes the parameters (xmid, scal and maxc) that characterise the position and shape of the sigmoidal curve. The dose-response curve is then used to fit the drug mapping function that governs the drug effect on intracellular signalling (see Figure S4). When a drug is available with a specific concentration in the surroundings of a cell, the cell response (viability) is looked up in the dose-response curve according to the available drug concentration. The dose-response curve is used to convert drug concentrations to cell viabilities and then, the corresponding drug target node is inhibited with a probability of $1 - cellviability$.

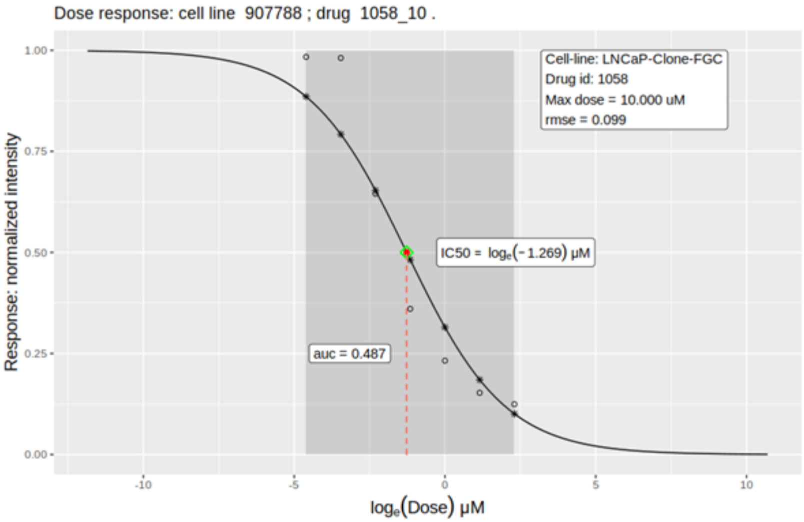


**Supplementary Fig S4 Drug-response curve for Pictilisib in LNCaP cell line fitted to the linear fixed effect model [15] with the *R* package *gdscIC50*.**

#### S4.3 Usage of the simulation setup tool

As part of PhysiBoSS 2.0, we created a simulation setup tool that can be accessed on GitHub: https://github.com/PhysiBoSS/PhysiBoSS/blob/master/addons/drugsims/physiboss_drugsim.py

With the help of this tool, all files and folders that are necessary to perform drug simulations are set up.

The setup tool enables the choice of:

- The generic project that contains the personalised Boolean model files. The project used in this work is "prostate" which was set up specifically to showcase PhysiBoSS 2.0 and can be found on GitHub: https://github.com/PhysiBoSS/PhysiBoSS/tree/master/sample_projects_intracellular/boolean/prostate.
- The cell line that is used for the drug simulations which in our case was LNCaP.

- The drugs that are used for the drug simulations which in our case were "Ipatasertib, Afatinib, Ulixertinib, Luminespib, Selumetinib, Pictilisib".
- The proportion of drug-resistant cells is for example "0.2, 0.8". If no drug resistances are desired, choose "0".
- The simulation mode can be either "single", "double" or "both". In the "single" mode drug treatments are applied only separately while in the "double" mode every possible pair of drugs is combined, the "both" mode does both single and double simulations. In this work we used the simulation mode "both" to obtain single and double simulations.
- The drug concentrations to be tested can either be specified with IC values or in uM. In our case we used: IC10 IC30 IC50 IC70 IC90. Which is also the default option.
- The specification of the number of replicates. In our case, we replicated each simulation 10 times.
- The cluster option creates a bash script to run the drug simulation with the SLURM workload manager.

Help for using the tool can be obtained with the following command:

```
python3 addons/drugsims/physiboss_drugsim.py --help
```

To obtain the files and folders for our showcasing project use the tool the following way:

```
python3 addons/drugsims/physiboss_drugsim.py -p prostate --cell
```

#### S4.4 LNCaP-specific Boolean model growth behaviour upon single drug administration

We simulated the prostate cell line LNCaP for 7 days with six drugs (IC10, IC30, IC50, IC70 and IC90) and without any drug in a three-dimensional space with PhysiBoSS 2.0. All simulations were replicated 10 times.

We studied the area under the curve (AUC) to compare growth behaviours upon drug administration and converted this into a growth index as explained in the main document. A growth index below zero indicates a reduction in growth upon drug treatment with respect to the untreated condition while a value over zero indicates an increase in growth.

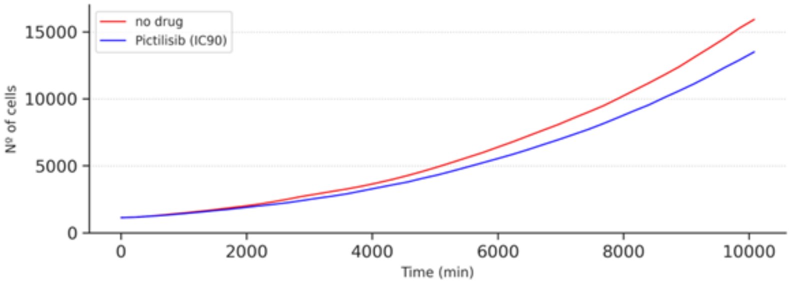


**Supplementary Fig S5 Growth curves for LNCaP with Pictilisib (blue) and no drug (red).** Pictilisib was given in a concentration corresponding to the IC90.

We performed a Kruskal-Wallis rank sum test to analyse the significance of growth behaviour changes for each single drug simulation, finding two out of six drugs significant. The two significant drugs are Ipatasertib and Pictilisib with $P \leq 0.0001$ (Table S2). Their growth indices are shown in Figures 2B and 2C (main text) in the first column/row indicated by "None".

#### S4.5 LNCaP-specific Boolean model growth behaviour upon double drug administration

We simulated pairs of drugs with the same simulation setup as in the previous section (Figure S4). We performed a Kruskal-Wallis rank sum test on the results to analyse the significance of growth behaviour changes for each drug combination. Among the 15 tested drug combinations, we found 12 significant combinations (Table S2). The two most interesting combinations with a significance $P \leq 0.0001$ were Ipatasertib + Pictilisib and Luminespib + Pictilisib. The growth index heatmaps for these two drug combinations are shown in Figures 2B and 2C in the main text and the full statistics for Pictisilib and Ipatasertib can be found in Figure S5.

All scripts and preprocessed data to obtain these figures can be found in https://github.com/PhysiBoSS/tools4PhysiBoSS-Drugs

**Table S2** P values for all single drugs and drug combinations obtained with a Kruskal-Wallis rank sum test. *: $P \leq 0.05$, **: $P \leq 0.01$, ***: $P \leq 0.001$, ****: $P \leq 0.0001$. 2 out of 6 single simulations are significant; 12 out of 15 drug combinations are significant.

| | | | | | | |
|---|---|---|---|---|---|---|
| Ipatasertib | 7.1e−5**** | | | | | |
| Afatinib | 1.4e−18**** | 8.7e−1 | | | | |
| Ulixertinib | 5.0e−19**** | 3.7e−2* | 9.2e−2 | | | |
| Luminespib | 1.9e−21**** | 7.3e−4*** | 7.2e−1 | 7.5e−1 | | |
| Selumetinib | 2.5e−19**** | 4.0e−3** | 2.5e−1 | 3.7e−1 | 6.2e−1 | |
| Pictilisib | 7.9e−28**** | 2.8e−16**** | 1.8e−15**** | 6.6e−27**** | 1.1e−19**** | 6.3e−7**** |
| | Ipatasertib | Afatinib | Ulixertinib | Luminespib | Selumetinib | Pictilisib |

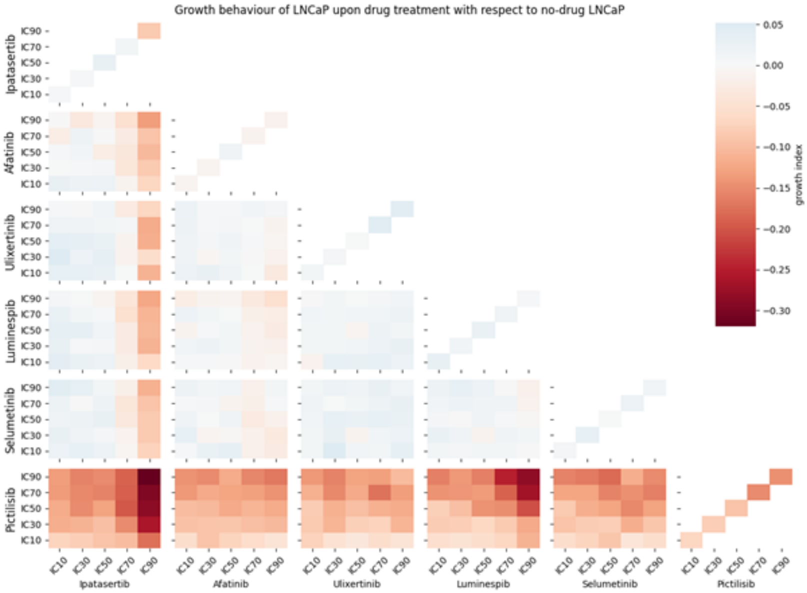


**Supplementary Fig S6 Growth indices for all single drug simulations (main diagonal) and double drug simulations (lower triangular matrix).** Every simulation was replicated 10 times. For each drug simulation, the growth index was calculated by taking the median AUC of the drug simulations and comparing it to the median without-drug AUC. White colour means no growth behaviour change upon drug administration, blue means the drug increased the growth and red means that the drug diminished the growth of the cells.

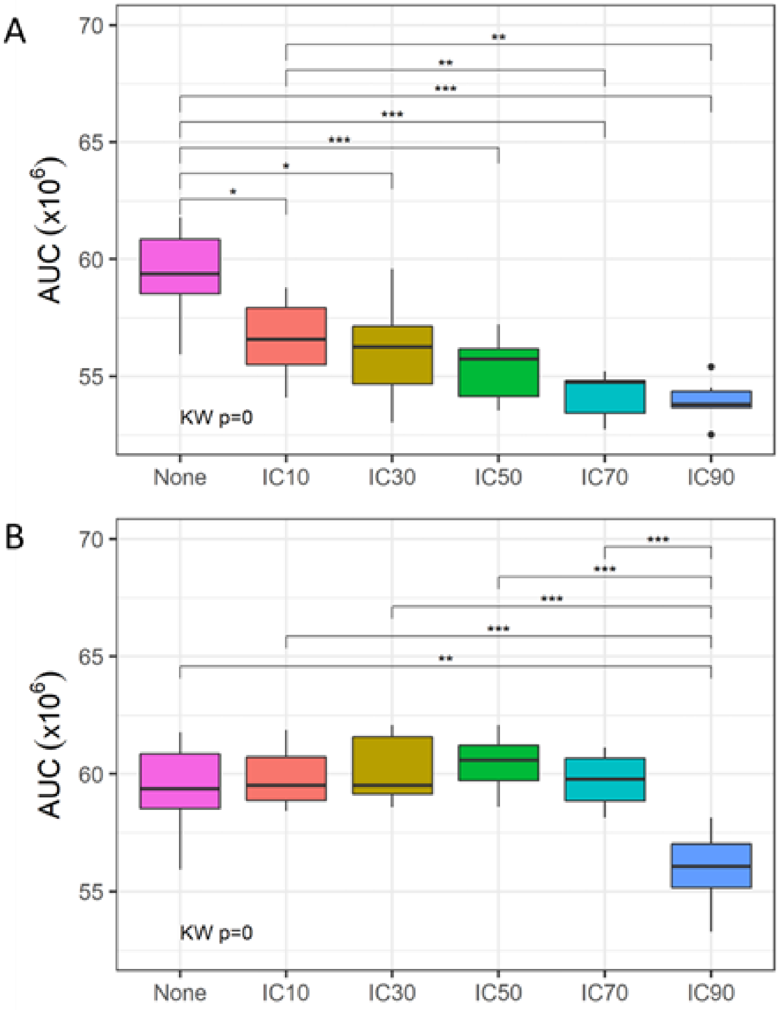


**Supplementary Fig S7 Dispersion of replicates for the untreated LNCaP and all drug concentrations with administration of Pictilisib (A) and Ipatasertib (B).** We performed a Kruskal-Wallis rank sum test for the complete set of concentrations and a pairwise Wilcoxon test to test for significance among pairs of concentrations adjusted with the Holm-Bonferroni method (*: $P \leq 0.05$, **: $P \leq 0.01$, ***: $P \leq 0.001$, ****: $P \leq 0.0001$). All statistics were done using the R stats package (https://rdrr.io/r/stats/stats-package.html).

#### S4.6 Double drug simulations synergy predictions

Synergistic drug combinations increase the anti-tumourigenic effect and reduce the likelihood of the development of drug resistances allowing for a reduction of drug dosages and, thus, minimising side-effects [16**?** ]. As we wanted to identify synergistic drug combinations, we calculated synergy scores using the Bliss independence model (Bliss, 1939) and the Combination index (CI) [17].

To calculate the Bliss reference model, we scaled the AUC data to values ranging from 0 to 1; 0 meaning no growth inhibition was obtained while 1 indicates the highest observed growth inhibition (Figure S6). Using Combination Indices, we found high synergy values for the two drug combinations Ipatasertib + Pictilisib and Luminespib + Pictilisib (Figure S7). Synergy values reach their maximum for Luminespib + Pictilisib with high drug concentration values of both compounds. Especially high drug concentrations of Luminespib drive the synergy value of the drug combination.

For Ipatasertib + Pictilisib, on the other hand, synergy values peak for IC values around IC50 and IC70 for both compounds. Comparably, high drug concentrations of Ipatasertib especially drive the synergy values. For both drug combinations, synergy values strongly decrease or even turn into slight antagonism with low drug concentrations for all drugs. Such varying combinatorial effects depending on the drug concentration have previously been observed as drug combinations can be both antagonistic and synergistic at the same time which can be described with the help of exposure-response surfaces [18]. These surfaces can help in choosing drug concentrations for combinatorial therapies.

In summary, using PhysiBoSS 2.0 as a drug simulation framework enables the performance of drug studies in cell populations. PhysiBoSS 2.0 can simulate the effect of the microenvironment in tumour cell populations, which plays a great role in cancer progression and the effectiveness of drugs. Therefore, it takes an important step towards the discovery of truly personalised drug treatments and effective drug synergies.

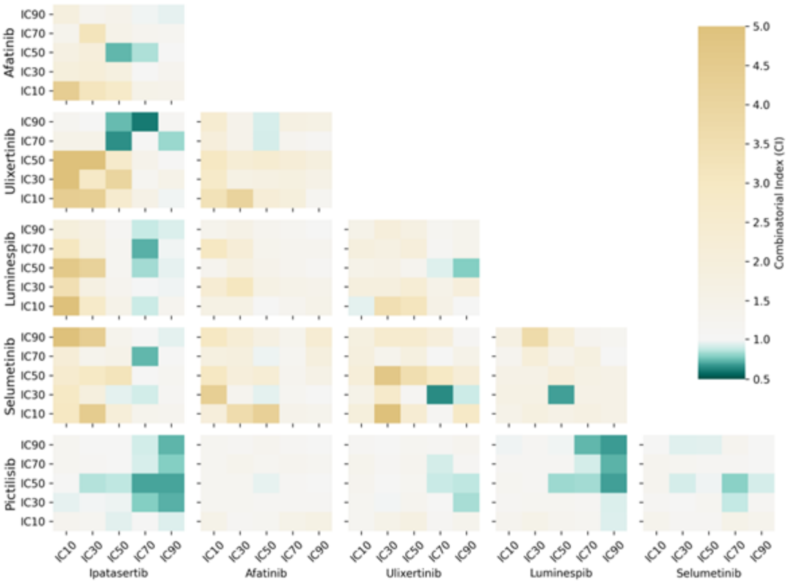


**Supplementary Fig S8 Complete heatmap of Bliss combination indices for all drug combinations.** White colour indicates an additive effect, green colour a synergistic effect, and yellow an antagonistic effect.

#### S4.7 Experimental validation of single simulations

To validate the obtained growth behaviour predictions for single drug administrations on LNCaP, we used experimental data from [12]. Real-time cell survival assay data using the RT-CES 96-well E-plate system was obtained for two drugs used in this study: Pictilisib and Selumetinib. Furthermore, this data was also available for the drug NMS-E973 which targets HSP90, similarly to Luminespib. However, validations based on drug-response data from two different compounds should be done with caution as drug-response relationships can differ across different compounds. Growth conditions for the cell survival assays resemble the conditions applied in the simulation with nutrients and growth factors in excess. Drugs were tested individually for the concentrations 3.3uM, 10uM and 30uM. A cell growth phase precedes the time of the drug administration which leads to varying cell indices at t=0. We calculated a centred cell index by normalising each drug condition to an initial value of 1.0 at t=0 to be able to compare across drug conditions. The cell index was normalised to a centred cell index to compare across drug conditions.

We performed simulations with corresponding drug concentrations while keeping IC values for the drugs on similar levels. For NMS-E973 we chose the lowest drug concentration available (3.3 uM) which corresponds to the IC92 of Luminespib (Figure S10). Accordingly, we chose a drug treatment of 10uM for Pictilisib as it corresponds to its IC90 (Figure S9). To fit the IC values also for Selumetinib we chose the highest drug concentration available (30uM) which corresponds to an IC16 (Figure S11). Unfortunately, IC values could not be matched up for all three drugs (Table S3). We then calculated the AUCs for the growth curves of the experimental data and performed simulations to determine the growth indices.

Table S3 shows that simulations consistently underestimate the effect of drugs on growth changes. But considering that, the trend of the three validated drugs is the same in the simulations and the experiments: Pictilisib is the most effective drug in terms of growth reduction; Luminespib is barely effective; and Selumetinib shows some pro-tumorigenic behaviour.

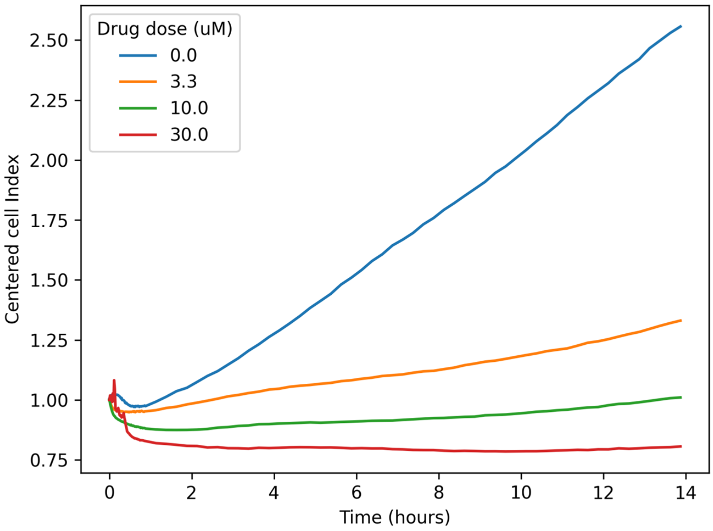


**Supplementary Fig S9 Dose-dependent changes of the centred cell index for the LNCaP cell line upon drug administration of Pictilisib at t=0.**

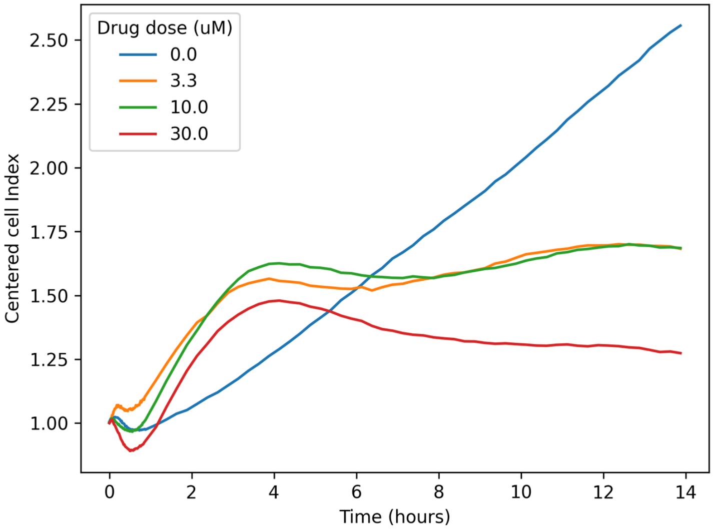


**Supplementary Fig S10 Dose-dependent changes of the centred cell index for the LNCaP cell line upon drug administration of NMS-E97 at t=0.** White colour indicates an additive effect, green colour a synergistic effect, and yellow an antagonistic effect.

**Table S3** Growth indices of drug simulations and experimental data for Pictilisib, Luminespib/NMS-E973 and Selumetinib. Pictilisib was given with a drug concentration of 10uM which corresponds to its IC90, NMS-E973 was given with a drug concentration 3.3uM which corresponds to the IC92 of Luminespib, Selumetinib was given with a concentration of 30uM which corresponds to its IC16.

| | **Pictilisib (10 uM ∼IC90)** | **Luminespib (∼IC92) / NMS-E973 (3.3 uM)** | **Selumetinib (30 uM ∼IC16)** |
|---|---|---|---|
| Drug simulation | -0.15 | -0.014 | 0.025 |
| Experimental data | -0.89 | -0.17 | 0.21 |

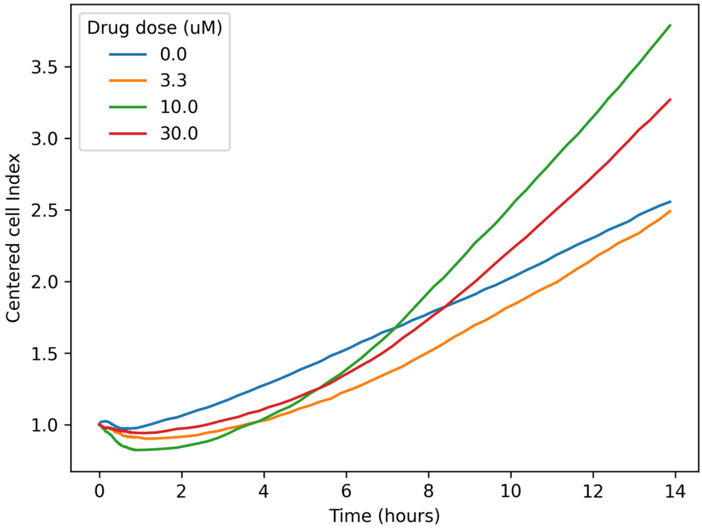


**Supplementary Fig S11 Dose-dependent changes of the centred cell index for the LNCaP cell line upon drug administration of Selumetinib at t=0.** White colour indicates an additive effect, green colour a synergistic effect, and yellow an antagonistic effect.

#### S4.8 Extended description of the heterogeneity in drug screening studies.

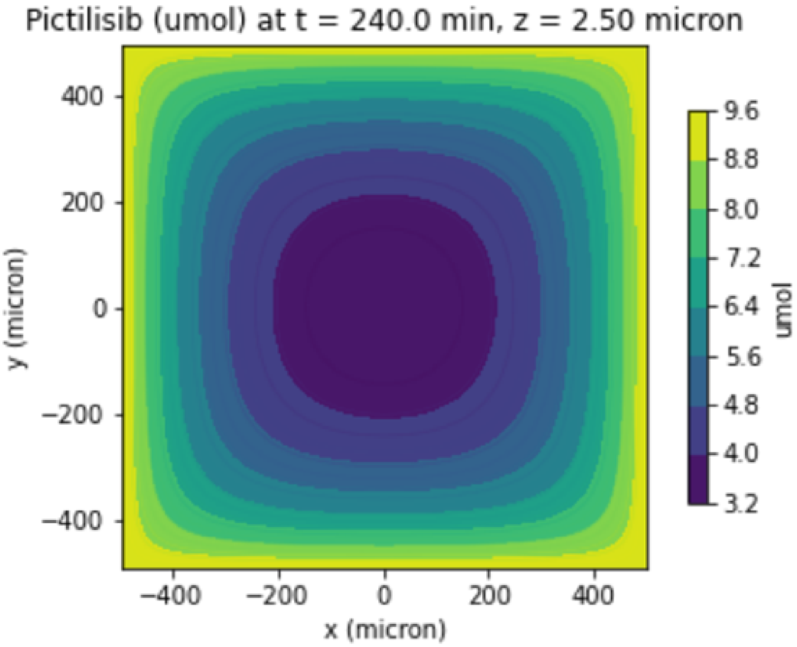


**Supplementary Fig S12 Pictilisib concentration throughout the computational domain along the z=0 plane at t=240 min.** The drug diffuses from the simulation boundaries inwards. Cells consume drugs and oxygen and, thus, reduce the substrate availability to other cells. Cells that are located in central regions of the tumour will have less substrate available than cells on boundary regions of the tumour and will be affected less by drugs.

*Genetic heterogeneity of the tumour*

PhysiBoSS 2.0 can also include heterogeneity caused by different genetic backgrounds or functional mutations. Increased mutation rates are very common in cancer, cause a high genetic heterogeneity among cells in the tumour and is one of the main causes of drug resistance in cancer treatments [19]. Drug resistances among tumour populations affect the treatment success of single and combinatorial therapies and are a harbinger of tumour relapse.

PhysiBoSS 2.0 enables to set-up of different cell strains, resistant or not to a given drug, and specifies their proportion among the cell population by using a simple python script (Section S4.3). While in the drug-sensitive cell strain, the drug leads to an inhibition of the corresponding node in the Boolean model, in the drug-resistant cell strain the node activity state remains unchanged (Figure S13).

By combining the microenvironment and genetic heterogeneities, PhysiBoSS 2.0 is an ideal tool to study drug resistance and tumour relapses in realistically complex cell populations. In this sense, future developments will be targeted towards the integration of cell-specific mutation rates of the nodes of the Boolean model to acquire drug resistances throughout the simulation.

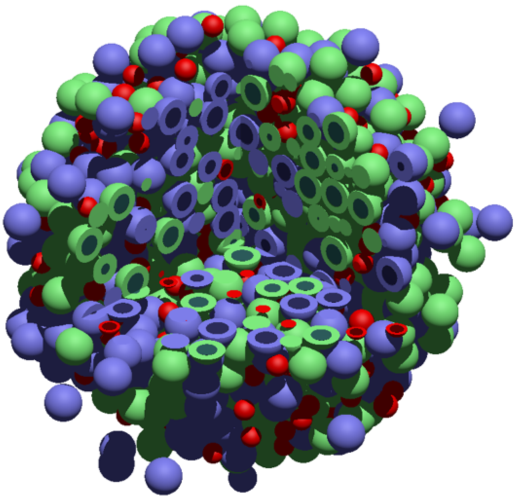

**Supplementary Fig S13 3D visualization of a cell population with two cell strains: green cells are sensitive to the drug while blue cells are resistant to the drug.** Red indicates apoptotic cells.

#### S4.9 Extended discussion of the multiscale simulations of drug treatments and combinations

This work presents the framework PhysiBoSS 2.0 to perform multi-scale drug simulations in cell populations. It enables the retrieval of personalized points of intervention, suitable drugs and their concentrations as well as drug synergies while capturing tumor heterogeneities. PhysiBoSS 2.0 is based on dose-response profiling data which enables the differentiation of therapeutic compounds. The personalization of cell lines in the form of the previously personalized Boolean models is enhanced with growth data from literature and, thus, represents specific characteristics of the cell lines better than the non-personalized model. Our framework is suitable for HPC systems and, therefore, supports high-throughput studies on large search spaces of therapeutic compounds and cell lines, as well as model exploration methods [20]. Furthermore, this work allows researchers to study different proportions of drug-resistant cell strains as well as other phenotypes typically triggered by cancer progression such as cell migration.

To showcase the use of the framework, we studied the prostate cell line LNCaP and 6 different drugs (Table 1). We identified two effective compounds, Pictilisib and Ipatasertib, as well as two synergistic drug combinations, Pictilisib and Ipatasertib and Pictilisib and Luminespib. The effectiveness of one of the identified compounds Pictilisib was validated experimentally (Table S3). Nevertheless, further experimental work is needed to validate other interesting compounds and synergistic drug combinations found by the simulations.

The limitations of our approach mainly originate in the data and models needed to personalize and fit the multiscale model. Firstly, our drug simulations consistently underestimate changes in growth behaviour obtained from experimental data. This might be due to the parameters used to map the activation of the Boolean Proliferation node to the cell cycle model used in the agents. Those parameters directly affect the impact of the Proliferation node on the division speed of cells. Implementing a higher impact of the Proliferation node on the cell division speed could reduce the observed differences. In order to do that, we could use model exploration methods to find parameters that would match experimental data [20]. Secondly, our approach is limited to the use of drugs with available dose-response data. We make use of the GDSC database [14] which currently comprises 138 anti-cancer compounds and their corresponding dose-response data. However, other compounds can also be integrated into the model if experimental dose-response data is available. The linear mixed effect model [15] used by GDSC and our framework needs to be fitted to the data to obtain a sigmoidal dose-response curve suitable for integrating into the drug simulations. This can be done with the scripts provided by the drug analysis repository as part of PhysiBoSS 2.0. Thirdly, to have truly personalized results, our framework requires a tailored Boolean model for the cell line to be studied: only cell lines with a previously tailored Boolean model can be studied or a generic Boolean model such as [21] needs to be personalized beforehand. This can be readily solved by using the PROFILE framework [11].

We used our framework to predict drug synergies for prostate cell lines. However, the approach can easily be applied to specific patients as well by using a patient-specific Boolean model and performing some minor parameter changes. The integration of pharmacokinetic parameters is another natural expansion. However, typical drug properties studied in pharmacokinetics (absorption, distribution, metabolism and excretion) correspond to the body as a whole [22] and have a difficult translation to our current cell-line-oriented setup. However, with regards to simulating patient-specific models, pharmacokinetics should be included into the model. Regarding drug transport into the agents, in the present framework, we only considered passive diffusion of drugs into the agents. The framework could be expanded to support other internalization mechanisms such as transmembrane mechanisms or endocytosis to further characterize specific drugs. Another interesting aspect could be the integration of cell-cell adhesion and motility parameters due to their role in cancer progression and the development of metastasis.

Our drug simulation framework PhysiBoSS 2.0 enables the performance of drug studies in cell populations. It considers the heterogeneity found in tumour cell populations and their microenvironment which plays a great role in cancer progression and the effectiveness of drugs. Therefore, it takes an important step towards the discovery of truly personalized drug treatments and effective drug synergies. We consider this framework a substantial contribution to reaching a truly personalized medicine and to helping improve treatments for cancer patients.

### S5 Comprehensive comparison of PhysiBoSS and PhysiBoSS 2.0

In this section, we describe the full details of the different implemented models used to reproduce the original simulation reported with PhysiBoSS 1.0. In order to reproduce the original results, we re-implement the spheroid TNF model in PhysiBoSS 2.0. The model was developed to investigate complex behaviours observed in cancer spheroids expose to different regimes of TNF supply, which consisted of the supply of TNF in pulses of different concentrations, duration and frequencies. The experimental result showed that continuous exposures to TNF resulted in cells becoming resistant to the effect of the cytokines, whereas short pulses of a certain frequency caused the reduction of the tumour[23].

#### S5.1 The TNF model version 1.0

In the original work [10] the authors implemented a model of a tumour spheroid in PhysiBoSS to mimic the experimental setup used to study the effect of TNF pulses in a tumour spheroid. In this model, each agent uses the Cell Fate Boolean network to rule the phenotype of the cells (e.g. proliferation, apoptosis) based on the amount of TNF present in the environment which is supplied following different regimes. The Cell Fate Boolean network is an adaptation of the model reported in a previous paper [24] which has a single input node which represents the signal triggered by the TNF when it is bound to the receptor, and it has three mutually exclusive readout nodes, $Proliferation$, *Apoptosis* and $NonACD$ that rule the behaviour of the cell. The diffusion and uptake of TNF are simulated using PhysiCell's core diffusion solver BioFVM. The mapping between the concentration, $TNF$ ($[TNF]$) in the nearest voxel of a cell and the corresponding input node of the Boolean model was done using a kinetic model and a step function. The kinetic model simulates how the TNF bound to its receptor in each individual cell agent and is given by the following equation:

$$\frac{[R^*_{tnf}]}{dt} = k^{tnf}_{bind}[TNF] - \lambda^{tnf}[R^*_{tnf}] \tag{1}$$

where $[R^*_{tnf}]$, $[TNF]$ and $[R^*]$ are the concentrations of the receptor, the TNF, and the TNF-TNFR complex, $k^{tnf}_{bind}$ is the TNF binding constant and $\lambda^{tnf}$ is the decay rate of the bonded TNF. Furthermore, the coupling between the receptor model and the Boolean network was done using a step function $H(x)$ which returns 1 when $x \geq \theta$ and 0 otherwise, for a given threshold $\theta$, $x$ correspond to $[R^*]$ (total TNF bound to the receptor) and $\theta$ and threshold parameter. The model can be found in the following repository https://github.com/gletort/PhysiBoSS and the details for running this version can be found in the following link https://github.com/gletort/PhysiBoSS/wiki/Example_Spheroid_TNF

#### S5.2 The TNF model version 2.0

The model proposed by [10] described in the previous section was re-implemented and extended to include a more detailed kinetic model of the TNF-receptor dynamics by [25] based on the known molecular biology of the receptor[26]. Specifically, the TNF receptor model includes the binding of TNF to the cell receptor at a given rate, the internalisation of the TNF-receptor complex and the recycling of the receptor. As in the previous version the kinetic model of the TNF receptor was implemented in each individual agent using the following set of differential equations:

$$\begin{aligned} \frac{[R_e]}{dt} &= -k_{bind}[R_e][TNF] + k_{recycle}[R^*_i] \\ \frac{[R^*_e]}{dt} &= k_{bind}[R_i][TNF] - k_{endo}[R^*_e] \\ \frac{[R^*_i]}{dt} &= k_{endo}[R^*_e] - k_{recycle}[R^*_i] \end{aligned} \tag{2}$$

where $[R]$, $[TNF]$ and $[R^*]$ are the concentrations of the receptor, the TNF, and the TNF-TNFR complex, respectively, and where the TNF-TNFR complex $[R^*]$ can be in two states, in the cell membrane ($R^*_e$) or internalized ($R^*_i$). Moreover, $k_{bind}$, $k_{endo}$ and $k_{recycle}$ are the TNF binding rate to a cell receptor TNFR, the rate at which the complex TNF-TNFR is internalised and the rate at which the TNF is degraded and the receptor recycled, respectively. The model is integrated numerically using a backward Euler method and the time scale is the same time scale as the one used to solve the micro-environment diffusion equations. The coupling between the receptor model and the Boolean network was done using a step function $H(x)$ as in the previous version of the model, but in this case, $x$ corresponds to $[R^*_e]$ (the total TNF-TNFR complex in the membrane).

Figure S14 shows a representation of the multi-scale TNF model and the main differences between the two versions of the TNF receptor extracellular model. Besides the difference, the way in which both models are simulated is the same. In both cases, the receptor model is integrated using

For the implementation of the model in PhysiBoSS 2.0 we first created a new sample project called spheroid_TNF_v2, which is a derivative of the template_3D sample project, and then used the add-on interface to re-implement the model and replicate the results of TNF experiments. We created a `custom_module` file in our sample project that defines the interface between MaBoSS and PhysiCell variables, i.e. how a PhysiCell variable affects the Boolean model and how the readout nodes of the Boolean model affect the agent behaviour in the following steps. The model can be found in PhysiBoSS 2.0 as part of the sample projects contained in the main repository https://github.com/PhysiBoSS/PhysiBoSS together with the instructions to compile and run the simulations. With the scheme of the custom class completed, we started to replicate the TNF experiments from the original PhysiBoSS work using PhysiBoSS 2.0. We configured the cell and microenvironment properties in an XML configuration file to include the TNF with the same physicochemical parameters used by **(author?)** [10].

Since we wanted to replicate how the TNF pulses were done by using a similar approach to PhysiBoSS, we added three custom parameters in the XML configuration file to configure the times ($duration_add_tnf$) and frequencies ($time_add_tnf$) of injections and added these parameters to the main configuration file of the sample project to perform the injections of TNF whenever required. We also created a function to parse the PhysiBoSS initialisation file of cancer cells and set up the initial tissue disposition in a user-determined, and used the same initial file used in the simulations reported using PhysiBoSS 1.0. Finally, The first thing we modified in the spheroid_TNF_v2 sample project *Makefile*, so that it compiles the aforementioned two classes from MaBoSS and PhysiCell in the add-on that have dependencies with PhysiBoSS 2.0.

#### S5.3 Simulating heterogeneous multi-cellular spheroids and extracellular matrix

To showcase the use of PhysiBoSS 2.0 to study the role of genetic and phenotype heterogeneity in cell populations, such as the one found in tumours, and to further validate the results of our new tool, we present here the same experiments performed with PhysiBoSS [10]. In this model, a heterogeneous spheroid of cancer cells is initialised using different mutants of the TNF model. Specifically, we have studied different experiments combining the wild-type TNF model together with two mutant strains: cFLIP+ IKK+, which greatly promotes proliferation; and CASP3+ Cytc+, which causes the cells to go to Apoptosis. These experiments were done in an oxygen-limited regime with a starting proportion of 75% wild type and 25% mutant strains.

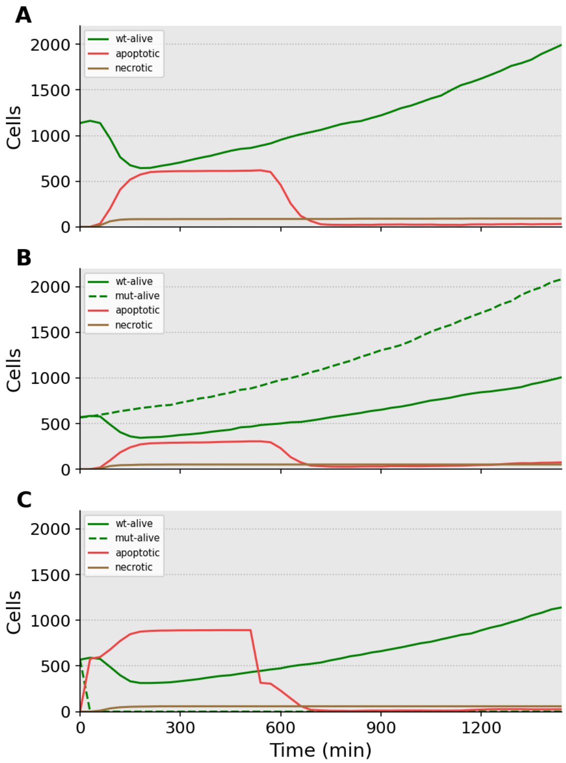


**Supplementary Fig S14 PhysiBoSS 2.0 allows for the study of TNF treatment response of heterogeneous multi-cellular spheroids in an oxygen-limited regime.** Wild-type TNF model (A) was mixed with two mutant strains, cFLIP+ IKK+ (B) and CASP3+ Cytc+ (C), and run with a starting proportion of 75% wild type and 25% mutant strains. These results reproduce the ones from Figure 5D of Supplementary File S2 in the PhysiBoSS paper.

The results from Figure S3 (cFLIP+ IKK+ shown in panel A and CASP3+ Cytc+ in panel B) reproduce the ones from Figure 5D of Supplementary File S2 in the original PhysiBoSS paper [10].

One remarkable improvement of PhysiBoSS 2.0 with respect to its original version is that now in PhysiBoSS 2.0 users can specify in the `XML` configuration file several cell definitions, each one using a different Boolean model. This was not that simple in PhysiBoSS 1.0 were having different Boolean models required tweaking internal parameters to force different nodes to have a given value, making this process more complex and rigid.

For completeness' sake, we also present an example in which cells can move across an extracellular matrix density, much like in PhysiBoSS' example_cells_with_ECM ( (https://github.com/sysbio-curie/PhysiBoSS/wiki/Example_Cells_With_ECM) and shown in Figure 2 of the [10].

This can be studied in a branch of the main code as this example uses beta features from PhysiCell: (https://github.com/sysbio-curie/PhysiBoSSv2/tree/example_physibossv2_ECM. Figure S4 shows the results of the experiment.

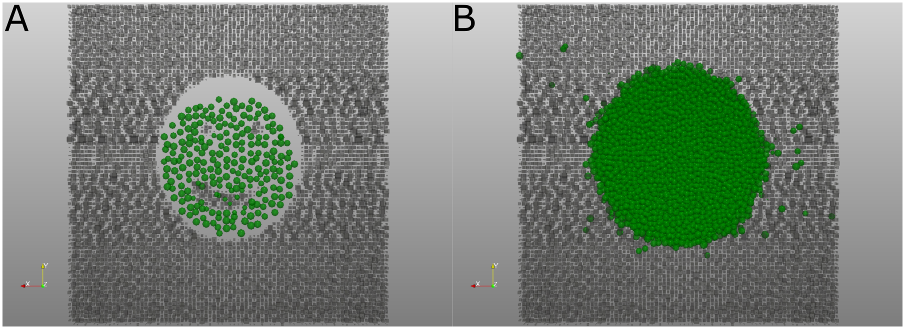


**Supplementary Fig S15 PhysiBoSS 2.0 allows for the study of cell migration and their interplay with the extracellular matrix.** PhysiBoSS 2.0 can integrate agents (green spheres) and ECM (grey boxes), as depicted at 0 hours (A) and 60 hours (B) of simulation. This example corresponds to the ECM example depicted in Figure 2 in the [10] paper and in PhysiBoSS' GitHub repository wiki.
